# One does not simply grow well: Performance of grassland plants in home and foreign soil and climate

**DOI:** 10.1101/2023.02.03.526963

**Authors:** Karoline H. Aares, Torunn Bockelie-Rosendahl, Ribha Priyadarshi, Francisco I. Pugnaire, Christian Schöb, Mohamed Alifriqui, Esteban Manrique, Laura H. Jaakola, Kari Anne Bråthen

**Author notes:** corresponding author: Karoline H. Aares [ ].

## Abstract

1. Plant-soil feedbacks (PSF) play a substantial role in determining plant performance in native and foreign habitats. Yet, PSF strength may be altered by climatic parameters, creating synergies. Here, we assess performance of alpine grassland species in their native and in foreign soils in an experiment including different climates.
2. Using soil and seeds sampled from six alpine grassland sites spreading in a gradient from Northern Africa to Northern Europe, we compared plant performance in home soil and in five foreign soils, as well as home and foreign climates (simulated temperature and photoperiod in growth chambers).
3. We found that despite a high variability in plant performance between sites, plants generally performed better in their home soil and home climate, than in foreign soil or in foreign climate. However, an interaction between soil and climate effect caused this better performance in home soil to occur only when in foreign climate. Similarly, performance improved in home climate only when plants were also placed in foreign soil.
4. Synthesis: In contrast to predictions from the literature, no benefit from growing in foreign soils are indicated. At least on the short term, climate change alone is not suggested to affect native grassland plant performance. However, when introduced to a habitat with a similar climate to their native habitat, plants may perform as well as in their native range, but when introduced both to a new soil and climate, plants will do poorly. This finding sheds light on the interactive effect of climate and soil origin, as well as the possible success of plant introductions under a changing climate.

## Introduction

The last decades have brought extensive research on plant-soil feedbacks (PSF) (Kulmatiski & Kardol 2008; Van Der Putten *et al*. 2013; Pugnaire *et al*. 2019), a process by which plants condition their soil environment, in turn affecting plant performance (Bever *et al*. 1997). Observed PSF are the sum of contrasting effects that co-occur simultaneously (Van Der Putten *et al*. 2013), and often are more negative than positive (Klironomos 2002; Bever 2003; Kulmatiski & Kardol 2008; Van Der Putten *et al*. 2016). Negative PSF are driven by species-specific soil pathogens that accumulate over time, limiting plant performance (Van Der Putten *et al*. 2016). Opposite, decomposers and soil mutualists drive positive PSF, improving plant performance (Bever *et al*. 2010; Van Der Putten *et al*. 2016; Palozzi & Lindo 2018). The prevalence of negative PSF in the literature may be a consequence of the methodological approaches typically used in PSF research, which normally apply monocultures of same versus other species to condition soil before assessing feedbacks, as plant growth in “home” versus “foreign” soil (Bever *et al*. 1997; Van Der Putten *et al*. 2013; Forero *et al*. 2019). While such research has been crucial in determining PSF drivers, it fails to predict PSF in natural ecosystems where more than one species is conditioning the soil (Baxendale *et al*. 2014; Kulmatiski *et al*. 2016; Guerrero-Ramírez *et al*. 2019).

In alpine grasslands, which typically have a high species richness (García-González 2008), dilution of soil pathogens may attenuate negative PSF (Maron *et al*. 2011; Schnitzer *et al*. 2011; Thakur *et al*. 2021). To understand PSF in such diverse communities, it is thus necessary to address PSF at the community level rather than at the single-species level (Kulmatiski *et al*. 2012; Guerrero-Ramírez *et al*. 2019; Forero *et al*. 2021; Forero *et al*. 2022). In addition, the gap between greenhouse-generated PSF and observed PSF in the field (Heinze *et al*. 2016; Forero *et al*. 2019) has further stressed the need for PSF experiments approaching field conditions (Kulmatiski & Kardol 2008; Van Der Putten *et al*. 2013; De Long *et al*. 2019b) by, for instance, minimizing pre-experimental soil treatments such as soil sterilization and inoculation (Brinkman *et al*. 2010; Diez *et al*. 2010; Fry *et al*. 2018).

Climatic parameters such as temperature (Wu 2011; Van Der Putten *et al*. 2016) and photoperiod (Sinclair 2003; Adams 2005) directly affect plant performance, as well as indirectly through interactions with PSF (Bennett & Klironomos 2018). Plants are adapted to their native climate and might be stressed under a foreign climate (Carlen *et al*. 1999; Joshi *et al*. 2001; Evert 2013). Additionally, climate impacts soil biota, thus influencing PSF (Frey 2008; Bennett & Klironomos 2018; De Long *et al*. 2019a). For instance, rising temperatures are expected to lead to more negative PSF due to increased pathogen growth (Burns *et al*. 2013; De Long *et al*. 2019a), coupled with decreased mycorrhizal activity (Mohan *et al*. 2014). How climatic parameters act together with PSF is increasingly important in a changing climate where species introductions across climatic barriers become more and more common (Van Der Putten *et al*. 2016; Pugnaire *et al*. 2019). Therefore, accounting for climatic effects in experimental PSF research can help build a more holistic understanding of current and future PSF.

The aim of this study was to determine whether alpine and tundra grassland species grow best in home or in foreign soil and whether the temperature and photoperiod of plants’ home climate may induce changes in performance compared with those of a foreign climate. We also aimed at identifying interactions between home and foreign soil and climate. Using soil and seeds from four alpine grasslands and two tundra grasslands from North Africa and Europe and simulating the alpine and tundra climates, we tested the hypotheses that (1) plants perform better in home than in foreign soil; (2) Plants perform better in their home climate than in a foreign climate. And finally (3) whether the effects of home versus foreign soils and climates interact.

## Methods

### Study sites

Soil and seed sampling sites were, from south to north, Oukaïmeden (Atlas, Morocco), Borreguiles area (Sierra Nevada, Spain), Néouvielle (Pyrenees, France), Flüela Pass (Davos, Switzerland), Varanger peninsula (Norway), and Adventdalen (Svalbard) (Table 1). The sites in the Atlas, Sierra Nevada, Pyrenees, and Alps are all alpine grasslands, while Varanger and Svalbard are respectively low and high Arctic tundra grasslands. Site location was selected so that environmental conditions were as similar as possible, and climatic variation among sites kept at a minimum. The Atlas and Sierra Nevada have an Oromediterranean climate, Pyrenees and the Alps are temperate and Varanger and Svalbard have a Polar climate (Geographic 2021). Sites differ in day length, day and night temperature, growth period length, and soil chemical composition (Table 1 and Figure 2).

**Table 1:** Climatic and geographic variation between study sites for soil and seed origins. In cases where the same reference is given for various sites, a place name search for the closest weather station to the respective sites has been used to obtain data for each site (essenc i.e., Oukaïmeden, Atlas, Morocco; Borreguiles, Sierra Nevada, Spain; Davos, Grisons, Switzerland; Vadsø, Varanger, Norway; and Longyearbyen, Svalbard). For the Pyrenees site, annual precipitation was calculated using the Gams index (Michalet *et al*. 2021) with data from the nearby weather station Aragnouet.

| Site | Climate regime | Coordinates | Altitude (MASL) | Yearly precipitation (mm) | Photoperiod mid-July (h/24h) | Growth period length (days) | Average daily min. and max. temperature in July (°C) |
| --- | --- | --- | --- | --- | --- | --- | --- |
| Atlas | Southern | 31°05N,<br>7°90E | 2700 | 603<sup>a</sup> | 12<sup>c</sup> | 120-149<sup>d</sup> | 17-34<sup>f</sup> |
| Sierra Nevada | Southern | 37°08N,<br>3°39E | 2800 | 690<sup>b</sup> | 14.5<sup>c</sup> | 150-179<sup>d</sup> | 18-31<sup>f</sup> |
| Pyrenees | Southern | 42°61N,<br>1°47E | 2100 | 2000 | 15<sup>c</sup> | 210-239<sup>d</sup> | 13-23<sup>f</sup> |
| Alps | Southern | 46°75N,<br>9°95E | 2300 | 1451<sup>a</sup> | 15<sup>c</sup> | 90-119<sup>d</sup> | 10-13<sup>f</sup> |
| Varanger | Northern | 70°42N,<br>29°42E* | 18 | 553<sup>a</sup> | 24<sup>c</sup> | 90-119<sup>d</sup> | 9-17<sup>f</sup> |
| Svalbard | Northern | 78°10N,<br>16°04E | 30 | 535<sup>a</sup> | 24<sup>c</sup> | 50-60<sup>e</sup> | 2-3<sup>f</sup> |

\* Subsites in Varanger cover a larger area than the remaining sites by being spread out over several valleys, respectively: Risfjorddalen, Store Saltjern, Kibergselva, Vesterelva, Finnvika, Sandfjordelva.
<sup>a</sup> (en.climate-data.org 2021), <sup>b</sup> (Oliva 2011), <sup>c</sup> (timeanddate.com 2020), <sup>d</sup> (FAO-GAEZ 2012), <sup>e</sup> (Karlsen 2014), <sup>f</sup> (MeteoBlue 2022)

### Soil and seed sampling design

Between July and September 2018, a hierarchical sampling design was used in each field site, where soil sample replicates were nested in subsites (Supplementary Material, Table S1). Subsites were located at least 100 meters apart from each other and replicates were located at least two meters apart. At each replicate, five-meter long transects were established and the aboveground plant community was described using the point intercept method to genus accuracy. Genus accuracy was considered adequate for this study since its field sites typically had only very few genera with more than one species present. Organic soil was sampled at each meter mark along the transect, to be pooled for each replicate. Soil was sampled down to 10 cm depth, and each replicate consisted of at least 2 dl of soil. All soil replicates were kept separate during the experiment.

In each sampling site, we collected seeds of abundant graminoids and forbs. Seeds of the same species were mixed for each site so that we ended up with one pool of seeds per species from each of the six sites. In October 2020, additional seeds were collected from Sierra Nevada and the Alps.

### Soil and seed preparation

With the cross-Europe site distribution of the present study, soil treatment for transport was unavoidable. We chose to dry the soil both for practical reasons and because we expected the effect of drying to be minimal (Blake *et al*. 2000; De Nobili *et al*. 2006), especially for grassland soils (Fierer *et al*. 2003). Furthermore, we assumed equal storage conditions for our grassland soils to be important (Martí *et al*. 2012; Meisner *et al*. 2021). Soil samples from all sites but Atlas were dried in a drying oven at 30°C for 48 hours, whereas Atlas soils were dried at 60°C for 48 hours. After drying, all soils were sieved to 2 mm mesh size. 10 cm diameter pots were prepared with a bottom layer of 0.5dl of Agra Perlite and a top layer of a homogenous mix of 1 dl Agra Perlite and 1 dl of soil. Pots were stored at room temperature during seed germination, which is described in detail in the Supplementary Material. Bar the abovementioned, no treatments were applied to the soils prior to planting in an attempt at keeping soils representative to field soils. In so doing, we arguably approximate a field-based experiment.

### Soil chemical composition analyses

From each soil sampling replicate, a sub-sample was analysed for elemental composition. Total carbon (C) and nitrogen (N) content were recorded using a LECO Truspec C/N analyser (St. Joseph, MI, USA). From this, carbon:nitrogen ratio (C.N) was calculated. Anion phosphate (PO_4_^3-^) was used as proxy for soil phosphorous (P) content, and was analysed by HPLC (Metrohm, HE, Switzerland) using concentrations in water extract (1:5 soil:water). Using soil N and soil P content, nitrogen:phosphorous ratio (N.P) was calculated. Other elements, such as calcium (Ca), sodium (Na), boron (B), potassium (K), magnesium (Mg), iron (Fe), zinc (Zn) and manganese (Mn) were determined after acid digestion with an inductively coupled plasma (ICP) emission spectrometer (ICAP 6500 DUO Thermo; Thermo Scientific, Wilmington, DE, USA). Soil chemical analyses were carried out at the CEBAS-CSIC ionomics lab (Murcia, Spain).

### Experimental design

We carried out two similar experiments, one in 2019 and one in 2021. Having failed germination of seeds from Sierra Nevada and the Alps in 2019, the first experiment only included plants from the Atlas, the Pyrenees, Varanger and Svalbard. To supplement our dataset, we re-sampled seeds from Sierra Nevada and the Alps in summer 2020 and used these in the second experiment. The protocols for the two experiments were identical, with standardized climatic conditions at the phytotron at UiT, The Arctic University of Norway. Hence, they are described together.

In this study, we considered the conditioning phase of PSF to have occurred in the field and hence be intact in the soil, allowing for the direct use of field soil to experimentally carry out the feedback phase.. Seedlings were planted in pots containing either home soil, being soil sampled at the same site as the seeds, or foreign soil, being soil sampled in each of the remaining sites (Figure 1). Seedlings were cleaned using distilled water and graminoids and forbs were planted together so that pots contained two individual plants, of different functional groups (See Table 2 for species list). Exceptions were plants from Pyrenees and Svalbard, where only one functional group was included due to germination failure. These differences were accounted for in statistical tests. Pots were kept on trays covered with transparent plastic sheets, which were gradually removed along the 11 weeks of the experiment. Plants were watered as needed. During the first three weeks plants were watered with distilled water. Afterward tap water was used.

**Figure 1.**
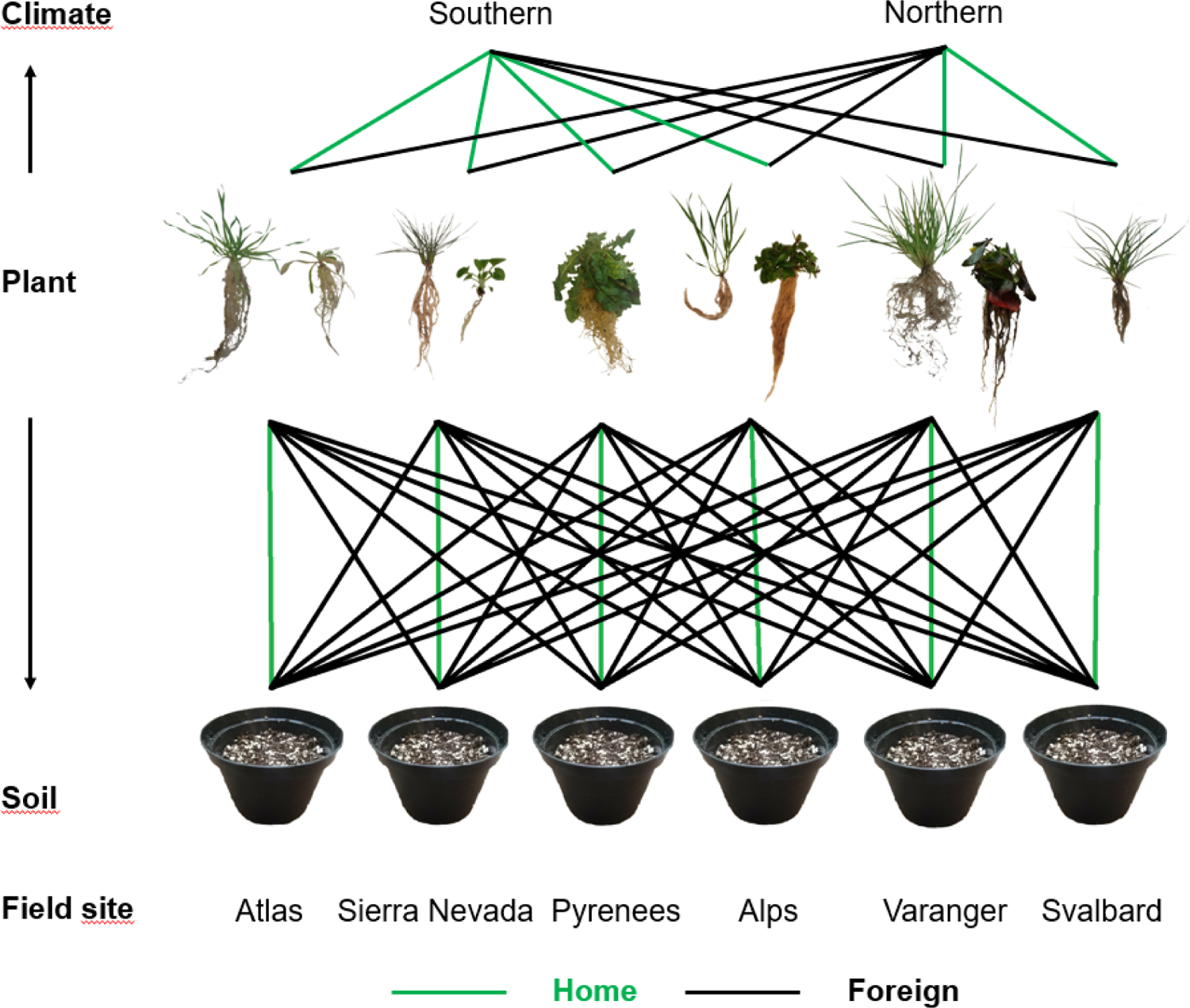
Planting design with seedlings and soil from six sites in Northern Africa and Europe where seedlings from each site were placed in all soils and both climates. Colour of line between plants and their designated soil and climate indicates whether it be home or foreign. Plants are illustrated using photographs of plants of the experiment after harvest and pots are illustrated with photographs of pots prior to planting.

**Table 2:** Plant species and their functional groups from each of the six sites used in the experiment.

| Plant origin | Functional group | Species scientific name |
| --- | --- | --- |
| **Atlas** | Forb | *Leontodon hispidus* (L.) |
|  | Graminoid | *Anthoxanthum odoratum* (L.) |
| **Sierra Nevada** | Forb | *Parnassia palustris* (L.) |
|  | Graminoid | *Nardus stricta* (L.) |
| **Pyrenees** | Forb | *Leontodon pyrenaicus* (L.) |
| **Alps** | Forb | *Cerastium alpinum* (L.) |
|  | Graminoid | *Phleum alpinum* (L.) |
| **Varanger** | Forb | *Rumex acetosa* (L.) |
|  | Graminoid | *Deschampsia cespitosa* (L.) |
| **Svalbard** | Graminoid | *Luzula confusa* (L.) |

The described setup was replicated in two growth chambers, each simulating one of two climate regimes (Figure 1) regarding photoperiod and temperature. In terms of photoperiod and temperature, sites within southern (Atlas, Sierra Nevada, Pyrenees and Alps), as well as northern (Varanger and Svalbard) sampling sites were relatively similar (Table 1). However, variation between southern and northern sampling sites was larger (Table 1). We assumed that climatic heterogeneity within these two groups would have negligible effects on plant performance at this scale (Table 1), allowing for a “southern” climate to be home climate for plants from Atlas, Sierra Nevada, Pyrenees and Alps, and an “northern” climate to be home climate for plants from Varanger and Svalbard. Temperature and photoperiod for the two climates simulated Pyrenees as southern climate, and Varanger, as northern climate. The Southern climate had 15 hours of light and 9 hours of darkness, with 15°C during light periods, and 9°C during dark periods. The Northern climate had continuous light, with 12°C for 12 hours, and 9°C for 12 hours. Day and night temperature fluctuations in the growth chambers used were limited to 3 degrees and thus diverges slightly from the true diurnal temperature variations in the field (Table 1). Each treatment combination of plant origin, soil origin and climate regime was replicated eight times, except for the Atlas which was replicated five times (Supplementary Material, Figure S1). This setup yielded data from 517 pots.

### Trait measurements

After 11 weeks, plants had reached the size in which competition within pots could have commenced and they were thus harvested. Soil was gently washed away from the roots and graminoids and forbs were separated by disentangling roots. We measured above and belowground properties to describe plant performance. Root length (Comas *et al*. 2013), root volume, canopy height (Pérez-Harguindeguy *et al*. 2013; Abeli *et al*. 2015) and shoot number were measured for each single plant after harvest. Root length was measured as the average of the three longest roots down to millimetre accuracy. Root volume was calculated using root length and root diameter measurements (described in detail in Supplementary Material, Figure S2). Canopy height was measured as the average of the three longest leaves from crown to leaf tip when stretched out. Shoot number was measured differently between functional groups because of the phenological differences in aboveground structuring between the graminoids and forbs in this study. “Shoot number” of forbs was in fact measured as number of leaves per plant (Funk *et al*. 2007; Li *et al*. 2015), while shoot number of graminoids was measured as number of aboveground shoots per plant (Soudzilovskaia *et al*. 2013). These were grouped together as one variable since both similarly describe aboveground growth investment. Above and belowground plant parts were separated, dried at 60°C for 48 hours and weighed to the mg (De Vries & Bardgett 2016). Above- and belowground mass was summed to get total dry mass per plant. The root-mass fraction (RMF) was calculated as the belowground dry mass divided by total plant mass (Poorter *et al*. 2012; Pérez-Harguindeguy *et al*. 2013) as a measure of biomass allocation. RMF was preferred over root-shoot ratio in this study because it yields constrained and arguably more comparable observations than root-shoot ratio, which can vary between very small and very large numbers. In pots where forbs and graminoids were planted together (i.e., plants from the Atlas, Sierra Nevada, the Alps and Varanger), all measured traits were averaged between plants from the same pot in order to avoid bias from root tangling within pots. Consequently, the final dataset included one observation per trait per pot.

### Data analyses

Data analyses were conducted using R 4.0.5 (R Core Team 2021). Non-metric multidimensional scaling (NMDS) with Bray-Curtis distance measures in the Vegan package was used to categorize variation in soil chemical composition within and between sites (Borcard 2011). Root length, root volume, belowground biomass, canopy height, shoot number and aboveground biomass were scaled using the scale function from base R (Becker 1988) and then added up for each pot. The resulting variable was named the Plant Performance Index (PPI), which rates overall plant performance in each pot relative to the pot mean in the dataset. Thus, positively deviating PPI suggest higher overall performance. To address effects of biomass allocation, RMF was analysed simultaneously with PPI. These two parameters were plotted in the NMDS to test for correlation with soil elements, determining which soil elements best explained variation in PPI and RMF.

We used linear mixed effect models in the Lme4 package (Bates *et al*. 2015) to compare PPI and RMF among the different treatments. PPI was log-transformed prior to linear modelling as it was right skewed (Mangiafico 2016). We built two separate models with PPI and RMF as respective response variables. Models fit were checked using diagnostic plots for a linear mixed model in the predictmeans package (Luo 2021). Models were built using the backward stepwise selection approach, starting with models including all predictors that may have affected PPI or RMF and followed by the removal of predictors which did not explain any variation. In line with our research questions, main predictors were home versus foreign soil and climate, as well as an interaction between the two. Additionally, we included number of plants per pot, content of selected soil elements and genus richness in field sites as cofactors. Number of plants per pot was included in the analyses to test whether having different species numbers affected the results. Soil elemental content was included to detect the relative importance of soil chemical quality to the remaining predictors on plant performance. Using the NMDS (Figure 2), soil parameters were selected based on extent of variation between sites and correlation to the model’s response variable (PPI or RMF). Among correlating parameters, the one with the largest variation was selected. Plant origin (each of the six study sites) and the hierarchical soil sampling design (site/subsite/replicate) were included as random factors, as differences in PPI and RMF between plant origins and from different soil replicates were of no interest to the research questions. Due to singularity of fit, only soil could be kept as random factor in the linear model using RMF and all other random factors were thus removed from that model. Backward elimination for linear mixed effects models in the LmerTest package (Kuznetsova *et al*. 2017) was used to detect predictors which did not significantly explain data variation. Such co-factors, when inessential to the research questions, were then removed from the linear models. No changes were however done to the random structure of the model using PPI as response variable.

## Results

### Soil chemical composition in soil sampling sites

Soil chemical composition varied between sites (Figure 2, Supplementary Material, Table S2). Notably, Atlas had the highest N content, while the Pyrenees had the highest P content. Sierra Nevada soils had relatively high levels of Na, while Svalbard soils had relatively high levels of B and K. Alps and Varanger soils had a relatively large variation in soil chemical composition. The soil parameters that displayed the largest variation between sites were soil Ca, N, C, N:P, C.N, B, K and Fe. Notably, soil P and soil C.N had strong correlations with PPI and RMF respectively and were thus selected as fixed factors for the linear model Additionally, soil C, K and Fe were included as fixed factors due to their large variation and that they didn’t correlate with other fixed factors in the model.

**Figure 2.**
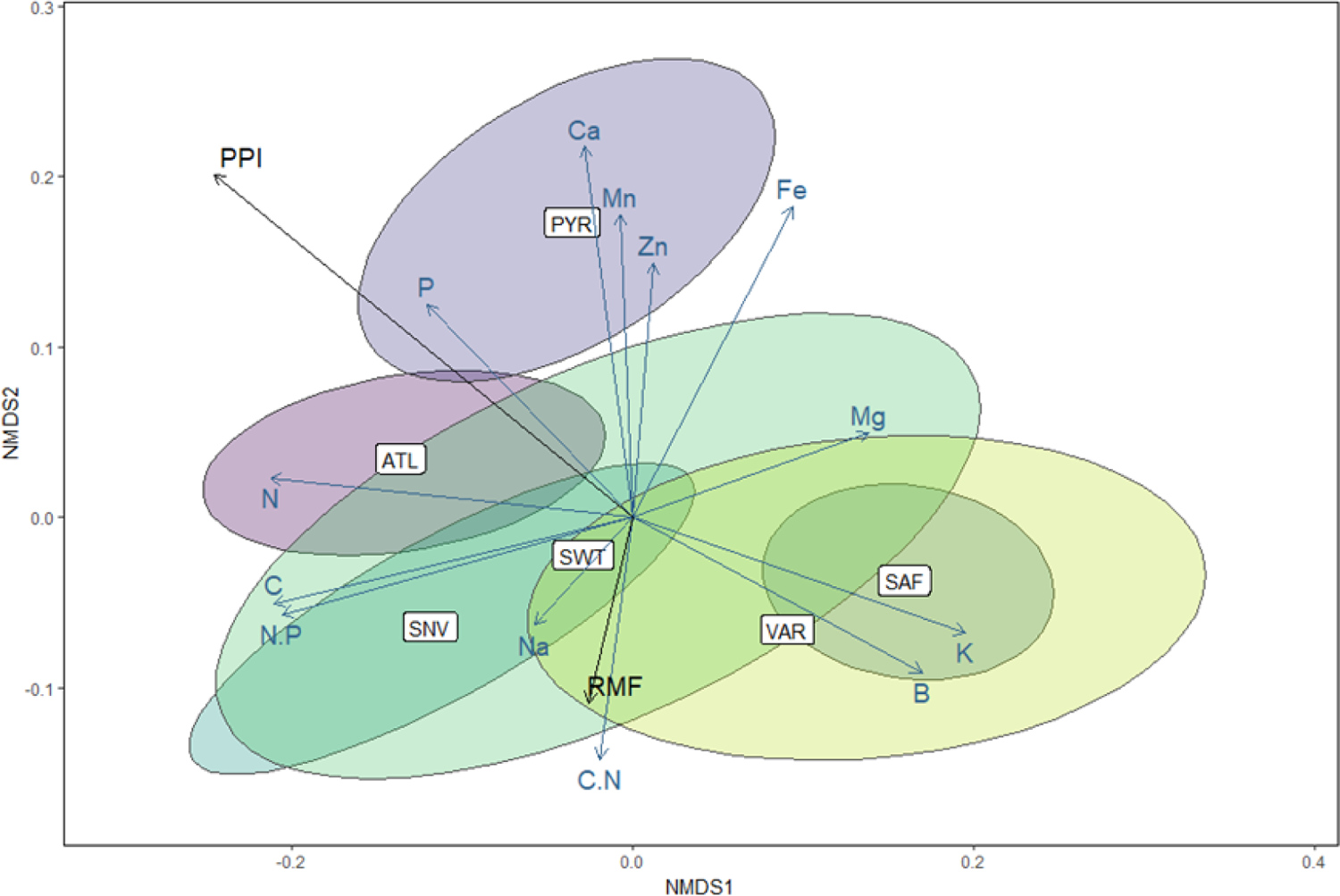
NMDS plot displaying variation between the six sites, Atlas (ATL), Sierra Nevada (SNV), Pyrenees (PYR), the Alps (SWT), Varanger (VAR) and Svalbard (SAF) regarding content of selected soil elements, nitrogen:phosphorous ratio (N:P) and carbon:nitrogen ratio (C:N). Ellipses are colour coded for each site and display location in the NMDS space where 95 percent of replicates for each site are found. Additionally, variation strength, correlation, and tendencies for highest content of the elements are displayed by blue arrows. Plant Performance Index (PPI) and Root-mass fraction (RMF) variation, correlation and tendencies towards higher values are displayed by black arrows. Arrow length indicates degree of variation and direction of arrows indicates where the highest levels are found, thus arrows pointing in similar directions show correlating parameters.

### Plant performance in soils

PPI varied considerably between the different soils (Figure 3). PPI was highest in Atlas soil for five out of six plant origins. All plant origins had the lowest PPI in Svalbard soils, and PPI in Varanger soils was also generally low. Performance responses to the remaining soils varied between plant origins (Figure 3). PPI was, overall, different in home and foreign soil (Supplementary Material, Figure S10), but this difference was highest for the plants grown in a foreign climate. Contrarily to PPI, RMF was very similar in the different soils (Supplementary Material, Figure S15).

**Figure 3.**
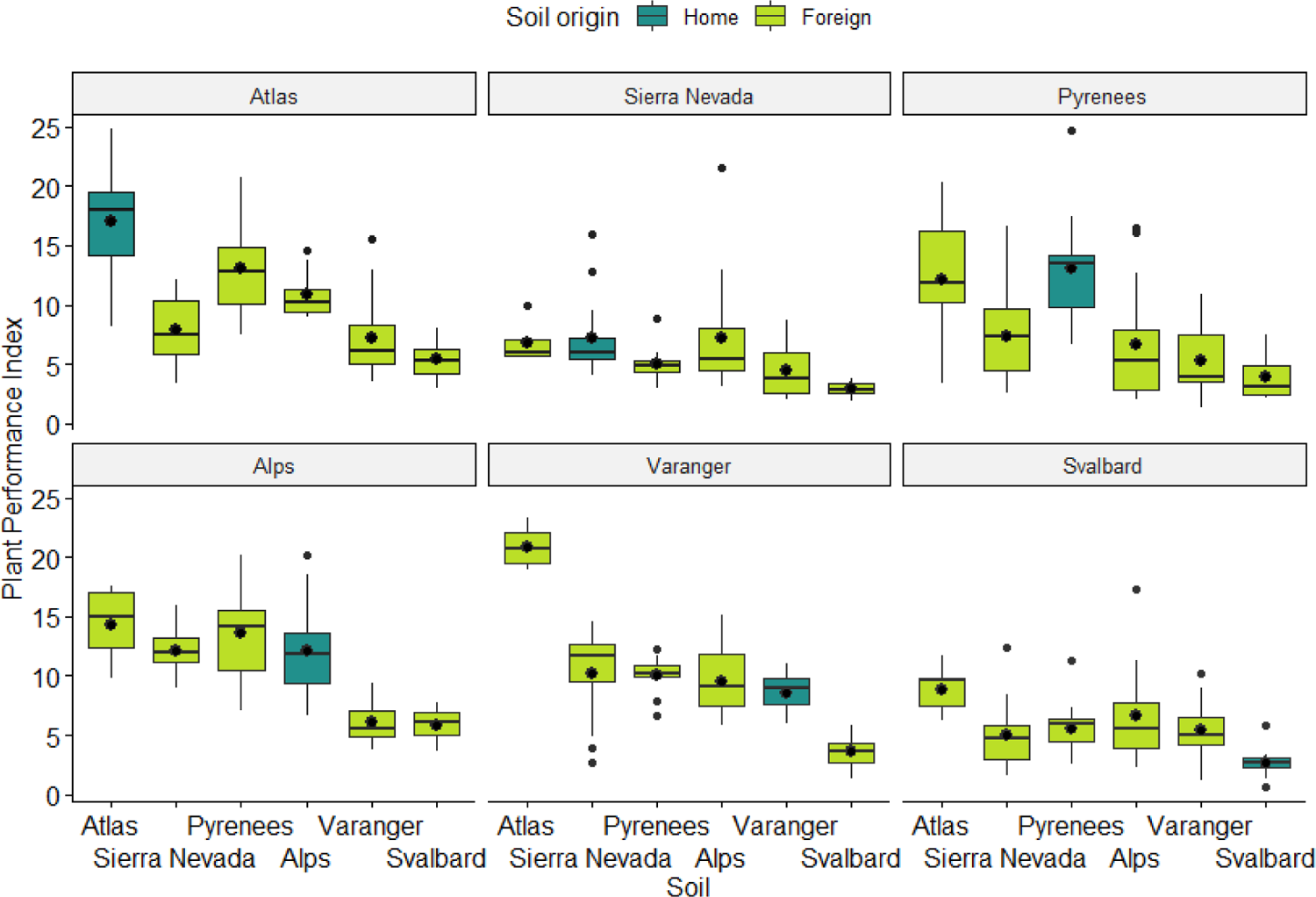
Plant Performance Index (PPI) in soils from each of the six sites, facetted by plant origin. Each box shows the mean (large black dot), median (middle horizontal line) and the lower and upper quartiles. Whiskers span the range of non-extreme values (less than 1.5 times the interquartile range), and the small dots show the outliers. Boxes displaying plants grown in their home soil are coloured blue, whereas boxes displaying plants grown in a foreign soil are coloured green.

### Plant performance in simulated climates

Plants from the southern climate regime (Atlas, Sierra Nevada, the Pyrenees and the Alps) had an overall slightly higher PPI in the southern than in the northern climate simulation (Figure 4 and Supplementary Material, Figure S11). Simultaneously, plants from the northern climate regime (Varanger and Svalbard) had a slight PPI increase in the northern climate simulation. This variation was highest for plants grown in a foreign soil. RMF did not vary between climate simulations (Supplementary Material, Figure S18 and S19).

**Figure 4.**
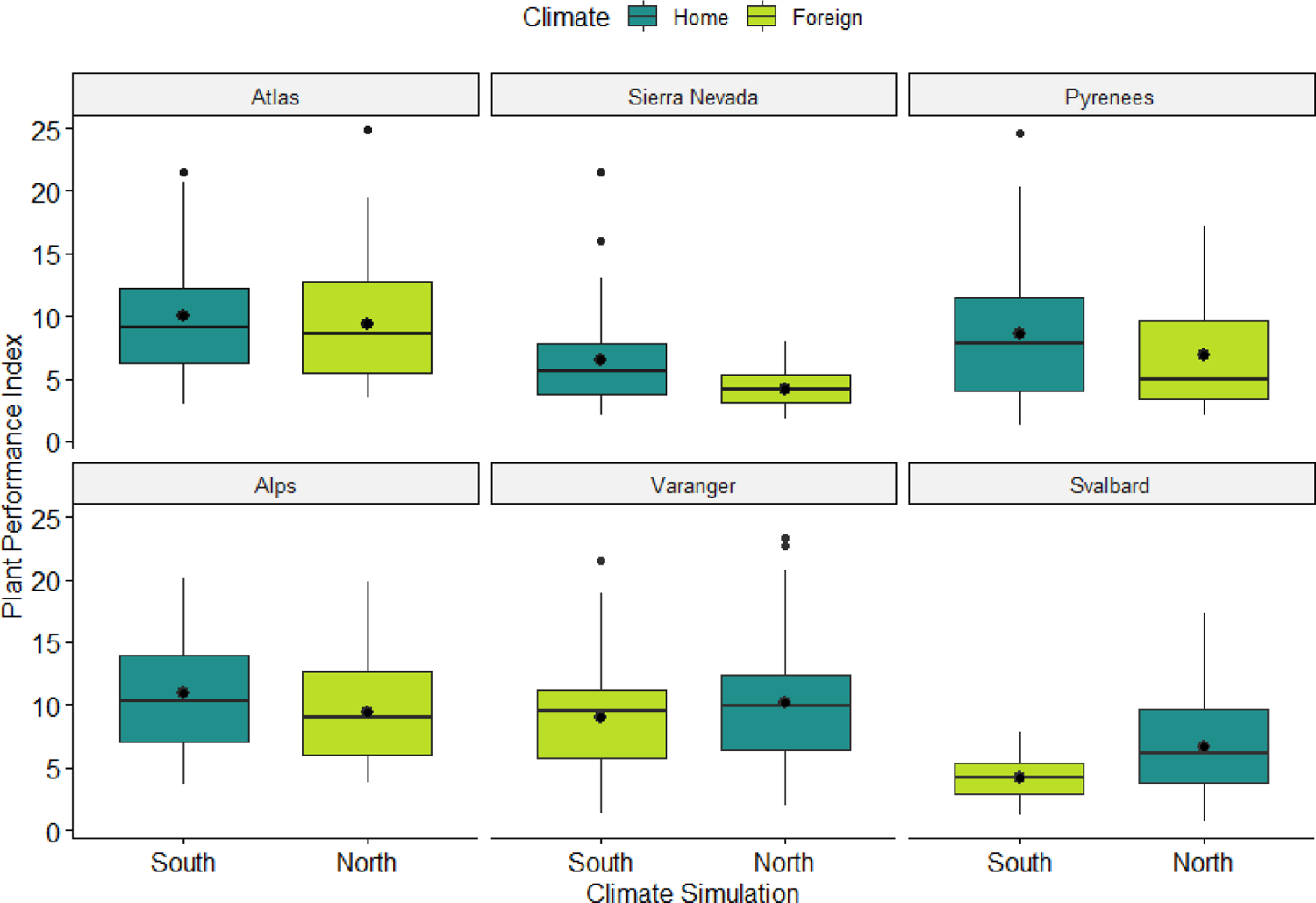
Plant Performance Index (PPI) in the two climate simulations, facetted by plant origin. Each box shows the mean (large black dot), median (middle horizontal line) and the lower and upper quartiles. Whiskers span the range of non-extreme values (less than 1.5 times the interquartile range), and the small dots show the outliers. Boxes displaying plants grown in their home climate are coloured blue, whereas boxes displaying plants grown in a foreign climate are coloured green.

### Linear mixed modelling

The final model for PPI included the main predictors; i.e., home versus foreign soil, home versus foreign climate and the interaction between these two (Figure 5a), as well as the additional predictors of soil P, soil C:N and soil K ( See Supplementary Material, Table S3 for details on model structure and output). There was no significant effect of the cofactor genus richness of grassland sites. There were no significant effects modelling RMF as response variable, (Supplementary Material, Table S4). In other words, RMF was unaffected by our treatments.

**Figure 5.**
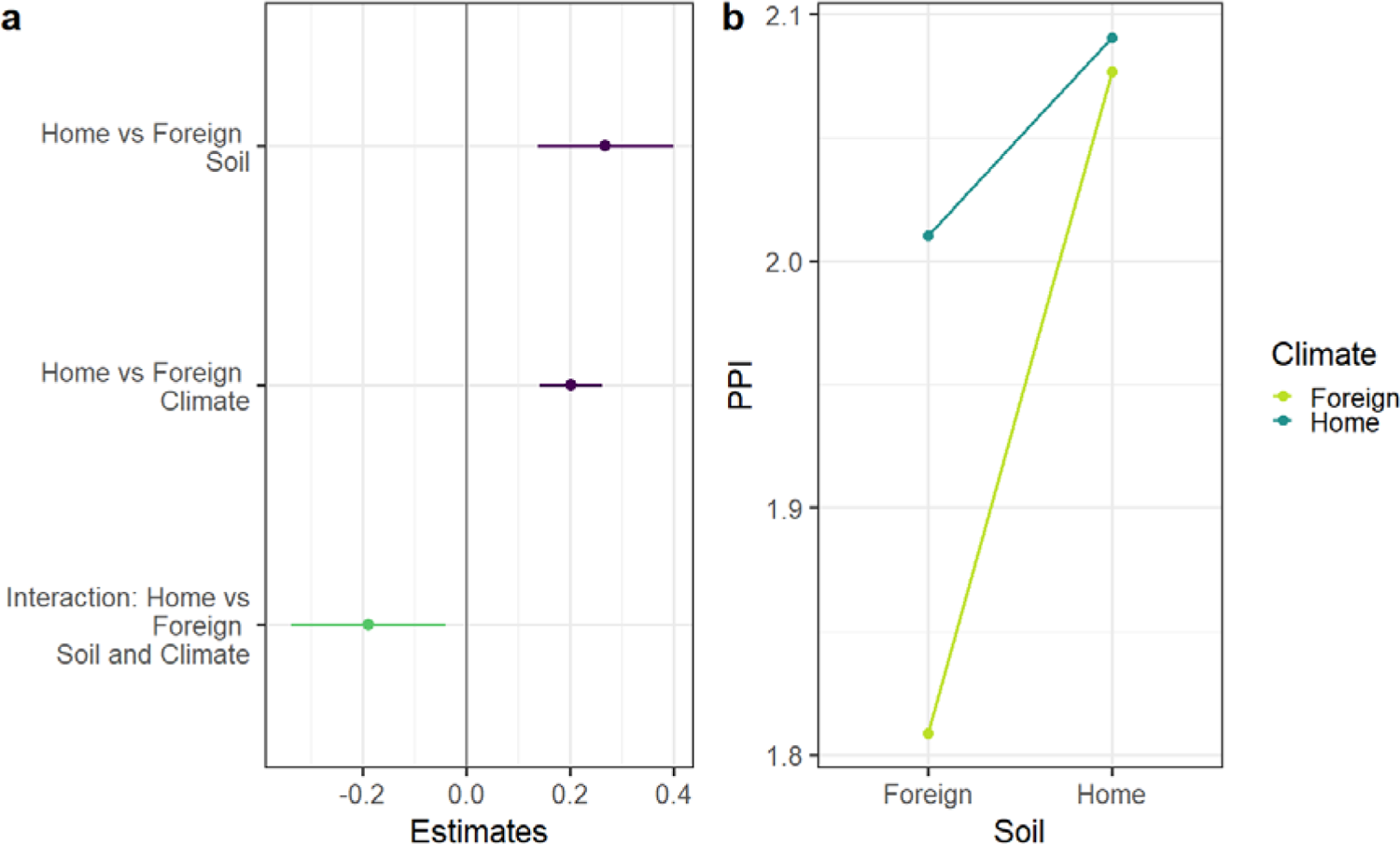
Linear mixed model of PPI (log transformed) as response variable and home versus foreign soil and climate as both additive and interactive predictors. Intercepts are foreign soil and foreign climate with home soil and home climate being one estimated unit increase from the intercept. Additional predictors are soil Phosphorous, soil C/N ratio, and soil potassium. Panels display the following: **(a)** Model estimates of main predictors. Positive estimates are purple and negative estimates are green, with error bars displaying mean ±CI of each estimate. **(b)** Interaction plot with predicted values for average PPI of plants in home and foreign soil and climate. Predictions for home climate are blue and predictions for foreign climate are green.

Plants had an estimated 31.1% higher PPI in home than in foreign soil. PPI was also 22.4% higher in home climate than in foreign climate. Both effects, soil, and climate, negatively interacted on PPI. PPI increase in home soil decreased by 21% when plants were grown in home climate (Figure 5b) and vice versa, PPI increase in home climate decreased by 21% when plants were grown in home soil. Ultimately, benefits from home soil were non-significant in home climate, and benefits from home climate were only observable in foreign soils. PPI was strongly enhanced with higher soil P. Increasing soil C:N and soil K had negative effects on PPI.

## Discussion

Our data show that grassland species perform better when growing in home soils and in home climate, aligning with research suggesting that environmental adaptation causes better performance in home environments (Joshi *et al*. 2001; Leimu & Fischer 2008; Bennington *et al*. 2012). Performance was enhanced in home compared to foreign soil in plants from four out of six sites, while one site had a neutral response, and plants from one site had a better performance in foreign than in home soils. There was a negative interaction between home soil and foreign climate effects, thus plants growing in a foreign climate could perform well as long as they grew in their home soil. Similarly, plants grown in a foreign soil would reach a high performance given that they were in home climate. However, when in both foreign soil and foreign climate, plants grew poorly. Although plant performance was strongly positively affected by soil P content and negatively affected by soil C:N and soil K, this did not superseeded the effects of soil and climate origin. Genus richness had no significant effect on PPI. Notably, all field sites had a genus richness exceeding the scale in which diversity effects on PSF is commonly studied (Kulmatiski *et al*. 2016; Guerrero-Ramírez *et al*. 2019; Forero *et al*. 2021). If plant performance and genus richness follow a positive saturating relationship (Weisser *et al*. 2017; Forero *et al*. 2021), it is likely that our systems were at the stage in which responses to increasing diversity are at a plateau.

RMF was similar under all treatments included in this study. In other words, biomass allocation was unaffected by soil origin or climate simulation, contrasting with the trends observed for PPI. According to the optimal partitioning theory, biomass allocation is an indicator of growth limitation, with plants investing more in organs increasing the uptake of the most limiting resource (Bloom *et al*. 1985; Bever *et al*. 1997; Shipley & Meziane 2002; Rehling *et al*. 2021). From our results on PPI, it is evident that soil origin, as well as soil phosphorous benefitted plant growth. The fact that RMF was not lower in home soil, nor with increasing nutrient content, is surprising, and in contrast with studies that reported a varying biomass allocation due to PSF (Veen *et al*. 2014; Xi *et al*. 2019), soil microbiology (Veresoglou *et al*. 2012; Borda *et al*. 2021), or soil chemical legacy (Delory *et al*. 2021). Our results however give no indication of such a plastic response to environmental variation, supporting instead the allometric partitioning theory, which predicts biomass allocation to be a function of plant size (Yang *et al*. 2009; Peng & Yang 2016; Liu *et al*. 2021).

Nutrient rich soils benefit plant performance (Gustafson & Casper 2004). The higher PPI in soils with higher P content confirms this, along with the tendency for a higher performance in soils that also had high soil N content, such as Atlas soil. Conversely, the low PPI in Varanger and Svalbard soil may be caused by their low P and N content, or the high levels of soil K and C:N. While soil K is an important element for photosynthesis, excess levels will limit Mg uptake (Tränkner *et al*. 2018), and slow down plant growth due to inhibition of crucial plant physiological functions (Xu *et al*. 2020). The higher C:N associated with Varanger, Svalbard and, to some extent, Sierra Nevada soils, limited plant performance, possibly due to low levels of accessible N caused by slow decomposition (Brust 2019). Importantly, growth in Svalbard soils was most likely limited also by the unintended drought in the experiment. These soils were highly inorganic and thus dried out frequently to moisture levels far below those typical for field conditions (Moriana Armendariz *et al*. 2021). Svalbard soils were kept in the model because their removal failed to change model estimates, but interpretations to be made on the Svalbard case are limited. Overall, the described responses to soil chemistry were general for the arguably most fertile soils, such as Atlas soils, as well as the less fertile soils, being Varanger (and Svalbard) soils. However, responses to the intermediately fertile soils from Sierra Nevada, Pyrenees and the Alps were more plant species-specific. This is an indication that on larger gradients of soil chemistry variation, chemical properties may be the stronger predictor of plant performance, but that PSF may become significant in chemically more similar soils (Gustafson & Casper 2004; in’t Zandt *et al*. 2019).

The higher performance in home soil compared to foreign soil suggests that enemy release had little importance on plant performance. Our study thus contradicts reports predicting that, in most cases, performance is better in foreign soil due to negative PSF (Klironomos 2002; Kulmatiski & Kardol 2008; Van Der Putten *et al*. 2016). we should point here, however, that plant performance is driven either by local adaptation (Joshi *et al*. 2001; Smith *et al*. 2012), positive PSF (Bever *et al*. 2010; Veen *et al*. 2019; Thakur *et al*. 2021) or both together (Van Nuland *et al*. 2016; Kirchhoff *et al*. 2019). With the methodological approximations to a field-based experiment, our study gives indications that, in the field, positive PSF, possibly along with local adaptation, is more influential to plant performance than what typical greenhouse experiments predict.

Being plants adapted to photoperiods and temperatures from their native habitat, foreign photoperiods (Marchand *et al*. 2004) and temperatures (Parmesan & Yohe 2003; Alexander *et al*. 2018) might prove disadvantageous for plant performance. Interestingly, more than half the species in this experiment have native populations in both climate regimes considered (Tackenberg 2019). Thus, our findings suggest that individual plants will grow best in their respective home climate although conspecific populations may be thriving in different climatic regimes. This is in accordance with literature describing plant ecotypes (Oleksyn *et al*. 1992; Liancourt *et al*. 2013), underlining the importance of individual origin for how well plants cope with different climatic conditions.

Our data suggest that climatic parameters affect PSF function, as described elsewhere (Van Der Putten *et al*. 2016; Duell *et al*. 2019; Pugnaire *et al*. 2019). The fact that plant performance reached a plateau when either soil or climate were “home” point to an upper limit to plant performance. In our experiment, benefits from home soil allowed plants to compensate for disadvantages of a foreign climate, and vice versa. Reports on interactive effects between climate and soil --defined either as home or foreign-- is scarce, although at least one study has documented the existence of such synergies (Macel *et al*. 2007). To fully understand the combined influence of soil and climate on plant performance requires a higher research effort, as the causation behind this interaction remains to be explored.

## Conclusion

Alpine grassland plants grow better in home than in foreign soils and climates. However, soil and climate negatively interact, so that when either soil or climate is home, benefits of the other being also home are negligible. If the described trends apply to the field, it implies that plants may grow well in foreign soils when the climatic conditions resemble that of their native habitat. Such a phenomenon is found in nature when niche tracking alpine plants establish at increasing altitudes (Alexander *et al*. 2015; Alexander *et al*. 2018). Another implication is that native soil buffers effect on climate change, so that native plants can perform well, in spite of climate change.

Our findings align with literature on local adaptation (Joshi *et al*. 2001; Leimu & Fischer 2008) and plant ecotypic variation (Oleksyn *et al*. 1992; Valladares *et al*. 2014) but challenge the view on PSF being predominantly negative (Klironomos 2002; Van Der Putten *et al*. 2016). It is possible that the methodological approach used in the present study has rendered results more in favour of positive PSF than the approach used by the bulk of PSF experiments. While this study does not contradict research on PSF at single species level, it gives new insights to how climate, local adaptation and PSF may shape diverse communities such as alpine grasslands.

## Acknowledgements

We thank all the driven researchers and technicians that contributed to this project, in particular Richard Michalet and Christian Kindler, who contributed substantially to field sampling, and Leidulf Lund at the phytotron at UiT, The Arctic University of Norway, as well as Mathilde Horaud who helped with the experiment. This project was funded by the Spanish Research Agency MCIN/AEI/10.13039/501100011033 (grant CGI/2017-84515-R) and supported by UiT, The Arctic University of Norway.

## Author Contributions

Karoline Helene Aares (KHA), Francisco I. Pugnaire (FIP) and Kari Anne Bråthen (KAB) conceived the ideas and designed methodology. KHA, FIP, Christian Schöb, Mohamed Alifriqui, Esteban Manrique and KAB were responsible for, and/or carried out fieldwork. KHA, Torunn Bockelie-Rosendahl, Ribha Priyadarshi, Laura Jaakola and KAB carried out the experiment and KHA analyzed the data. KHA, with supervision from KAB, led the writing of the manuscript, and all authors contributed to the writing of the manuscript. All authors have given their final approval for publication.

## Data Availability Statement

Data and code are available from the corresponding authors upon request and will be uploaded unto the repository of UiT, The Arctic University of Norway; UiT Open Research Data upon manuscript acceptance.

