## Supplementary Material for "One does not simply grow well: Performance of grassland plants in home and foreign soil and climate"

### Germination methods

The germination of seeds and the experiment itself took place in the phytotron at UiT – Arctic University of Norway, Tromsø. Seeds from Atlas, the Pyrenees, Varanger and Svalbard were sown in moist sand and placed in darkness with 0.5ºC for six weeks prior to the experiment, to promote germination. Seeds from Atlas and the Pyrenees were germinated in rooms with 18ºC and 15 hours of light per 24 hours. Svalbard and Varanger plants were put to germinate in rooms with 18ºC and continuous light. The lamps used for lighting in this experiment were Philips 58 40, having 58W and colour temperature 4000 K, with a maximum intensity of 200 mol. Seeds from Sierra Nevada and the Alps were put directly to germinate in moist sand under similar conditions as the Atlas and Pyrenees plants.

### Plant regime considerations

While observing the data (Figure S11), an ad-hoc hypothesis arrived that there might be differences in response to home versus foreign soil between plants originating from the southern sites (Atlas, Sierra Nevada, the Pyrenees and the Alps) and the northern sites (Varanger and Svalbard). Thus, “plant regime” (Either southern or northern plant origin) was added as predictor to the inclusive model. This trend could however not be statistically verified, and the experimental setup was not directed at addressing such latitudinal biogeography related hypotheses. An experiment studying growth or performance of conspecifics from sites across a latitudinal gradient could be suggested to research this further.

### Tables

Table S1: Sample sizes in the soil collection design. For each site sub-replicates are nested within replicates and replicates are nested within subsites.

| Site | Subsite | Transect/Replicate | Sub-replicate |
| --- | --- | --- | --- |
| Atlas Mountains | 6 | 5 | - |
| Sierra Nevada | 5 | 6 | - |
| Pyrenees | 5 | 6 | - |
| Alps | 5 | 6 | - |
| Varanger Peninsula | 6 | 3 | 2 |
| Svalbard | 6 | 3 | 2 |

Table S2: Mean and standard deviation (sd) for content of selected soil elements, nitrogen:phosphorous ratio (N.P) and carbon:nitrogen ratio (C:N). Unit is displayed under element symbol.

| **Variable** | **Measurement** | **Atlas** | **Sierra Nevada** | **Pyrenees** | **Alps** | **Varanger** | **Svalbard** |
| --- | --- | --- | --- | --- | --- | --- | --- |
| **B**  **(mg/kg)** | Mean | 10.369 | 8.605333 | 5.806667 | 6.311333 | 17.22 | 45.225 |
|  | sd | 2.045724 | 2.927037 | 2.016011 | 1.825148 | 10.86654 | 6.6335 |
| **C**  **(g/100g)** | Mean | 20.14333 | 16.766 | 9.134 | 12.62333 | 12.04611 | 6.323333 |
|  | sd | 4.694154 | 7.445644 | 3.163668 | 8.860525 | 13.06673 | 2.263862 |
| **Ca (g/100g)** | Mean | 0.826 | 0.135667 | 2.163667 | 0.257667 | 0.292778 | 0.281667 |
|  | sd | 0.184009 | 0.043445 | 0.748034 | 0.14474 | 0.365172 | 0.11888 |
| **Fe**  **(mg/kg)** | Mean | 21079.37 | 18922.54 | 31314.84 | 16094.43 | 18959.04 | 29312.58 |
|  | sd | 5514.231 | 5825.714 | 7207.403 | 10235.76 | 7823.758 | 3714.109 |
| **K**  **(g/100g)** | Mean | 0.379 | 0.505 | 0.526 | 0.631667 | 0.707222 | 0.892778 |
|  | sd | 0.080058 | 0.138109 | 0.121275 | 0.097134 | 0.426157 | 0.146199 |
| **Mg**  **(g/100g)** | Mean | 0.572333 | 0.302 | 0.414667 | 0.463333 | 0.561111 | 0.499444 |
|  | sd | 0.151423 | 0.090949 | 0.08161 | 0.308515 | 0.360439 | 0.085162 |
| **Mn**  **(mg/kg)** | Mean | 495.783 | 146.307 | 1864.466 | 275.113 | 252.2656 | 352.3528 |
|  | sd | 359.7255 | 56.49651 | 1569.037 | 283.4174 | 144.8204 | 94.19859 |
| **N**  **(g/100g)** | Mean | 1.580667 | 1.066333 | 0.791667 | 0.904 | 0.699444 | 0.398333 |
|  | sd | 0.354342 | 0.358507 | 0.307348 | 0.590491 | 0.652682 | 0.155232 |
| **Na**  **(g/100g)** | Mean | 0.032667 | 0.114667 | 0.028 | 0.037333 | 0.015 | 0.042778 |
|  | sd | 0.006397 | 0.037021 | 0.007144 | 0.012847 | 0.005145 | 0.010178 |
| **P**  **(g/100g)** | Mean | 0.105333 | 0.083 | 0.107 | 0.094667 | 0.073333 | 0.058889 |
|  | sd | 0.017167 | 0.025346 | 0.030978 | 0.026226 | 0.044721 | 0.010226 |
| **Zn**  **(mg/kg)** | Mean | 47.622 | 67.86767 | 79.642 | 40.82167 | 60.66889 | 64.78278 |
|  | sd | 12.37064 | 22.35691 | 21.48354 | 10.34303 | 61.67997 | 9.343364 |
| **C:N** | Mean | 12.73333 | 15.36667 | 11.93333 | 13.2 | 15.44444 | 16.05556 |
|  | sd | 1.460593 | 2.025413 | 1.837039 | 2.21904 | 4.355149 | 1.862074 |
| **N:P** | Mean | 15.36667 | 13.1 | 7.6 | 8.966667 | 8.444444 | 6.444444 |
|  | sd | 4.286976 | 3.356414 | 2.774266 | 4.270858 | 3.63354 | 1.822158 |

Table S3: Model output for Linear Mixed models using PPI as response variable. The inclusive model (left) includes main predictors home versus foreign soil and climate, and their interaction, along with co-factors hypothesised to affect the PPI. In the final model (right), co-factors with no significant variance are excluded. The intercepts are by default the first observation alphabetically, or numerically depending on the predictor. Thus, estimates compare PPI in home soil and climate to foreign soil and climate, southern plant regime to northern plant regime, two to one plant per pot, and estimates increasing species richness as well as soil compound contents. For the interactions it further compares PPI when grown in home soil and climate to PPI when either or both is “increased” to foreign, and PPI when grown in home soil and plant regime being North, to when either or both is “increased” to foreign, or South respectively.

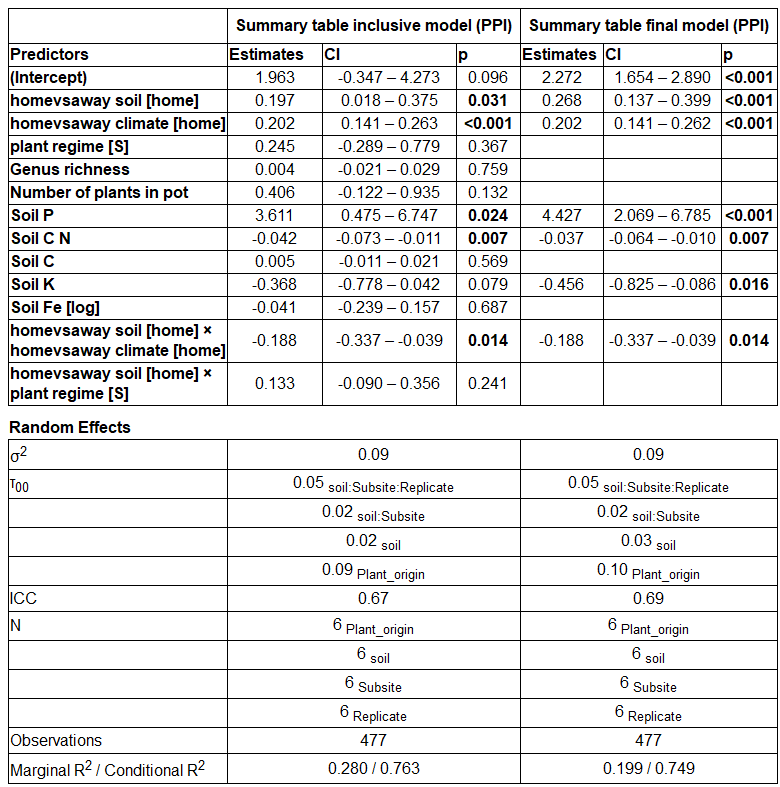

Table S4: Model output for a Linear Mixed model using RMF as response variable. The intercepts are by default the first observation alphabetically, or numerically depending on the predictor. Thus, estimates compare RMF in home soil and climate to foreign soil and climate, southern plant regime to northern plant regime, two to one plant per pot, and estimates increasing species richness as well as soil compound contents. For the interactions it further compares RMF when grown in home soil and climate to RMF when either or both is “increased” to foreign, and RMF when grown in home soil and plant regime being North, to when either or both is “increased” to foreign, or South respectively.

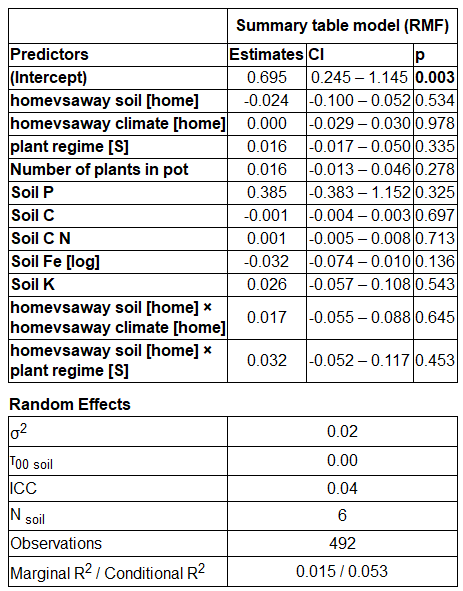

### Figures

|  | Climate | | | | | | |
| --- | --- | --- | --- | --- | --- | --- | --- |
| Soil |  | Home | | | Foreign | | |
|  | Home | ATL | T | | ATL | P | |
|  |  | ATL (5) | | | ATL (5) | | |
|  |  | SNV | T | | SNV | | P |
|  |  | SNV (8) | | | SNV (8) | | |
|  |  | PYR | T | | PYR | P | |
|  |  | PYR (8) | | | PYR 8) | | |
|  |  | SWT | T | | SWT | | P |
|  |  | SWT (9) | | | SWT (8) | | |
|  |  | VAR | P | | VAR | T | |
|  |  | VAR (8) | | | VAR (8) | | |
|  |  | SAF | P | | SAF | T | |
|  |  | SAF (8) | | | SAF (7) | | |
|  | Foreign | ATL | T | | ATL | P | |
|  |  | SNV (4), PYR (4), SWT (3), VAR (5), SAF (5) | | | SNV(0), PYR (5), SWT (1), VAR (2), SAF (3) | | |
|  |  | SNV | | T | SNV | | P |
|  |  | ATL (8), PYR (8), SWT (8), VAR (8), SAF (7) | | | ATL (8), PYR (8), SWT (8), VAR (6), SAF (8) | | |
|  |  | PYR | T | | PYR | P | |
|  |  | ATL(8), SNV (8), SWT (8), VAR (7), SAF (8) | | | ATL (8), SNV (8), SWT (8), VAR (8), SAF (8) | | |
|  |  | SWT | | T | SWT | | P |
|  |  | ATL (8), SNV (8), PYR (9), VAR (8), SAF (8) | | | ATL (7), SNV (8), PYR (8), VAR (8), SAF (8) | | |
|  |  | VAR | P | | VAR | T | |
|  |  | ATL (8), SNV (8), PYR (8), SWT (8), SAF (8) | | | ATL (8), SNV (8), PYR (7), SWT (8), SAF (8) | | |
|  |  | SAF | P | | SAF | T | |
|  |  | ATL (8), SNV (8), PYR (8), SWT (8), VAR (8) | | | ATL (8), SNV (8), PYR (8), SWT (8), VAR (8) | | |

| Soil site | Plant origin | Climate Simulation |
| --- | --- | --- |
| ATL – Atlas Mountains  SNV – Sierra Nevada  PYR – Pyrenees | ATL – Atlas Mountains  SNV – Sierra Nevada  PYR – Pyrenees | T – Temperate  P - Polar |
| SWT – Alps  VAR – Varanger Peninsula | SWT – Alps  VAR – Varanger Peninsula |  |
| SAF - Svalbard | SAF - Svalbard |  |

Figure S1: Planting design using soil and seedlings of six origins, and two simulated climates. Seedlings are planted in respectively all soil types, where “home” is soil originating in the same site as the seedlings, and “foreign” is soil originating from any of the other five sites. This setup is replicated in two simulated climates, a Temperate climate, and a Polar climate. The Temperate climate simulation has 15 hours of light and 9 hours of darkness per 24 hours, and temperatures is 15^o^C in light periods, and 9^o^C in dark period. The Polar climate simulation has 24 hours of light per 24 hours, where temperature is 12^o^C for 12 hours, and 9^o^C for 12 hours, per 24 hours. For Atlas, Sierra Nevada, Pyrenees and Alps plants, home climate is set as the Temperate climate simulation. For Varanger and Svalbard plants, home climate is set as the Polar climate simulation. Number of replicates for each combination of plant origin – soil origin – climate simulation is displayed in parentheses behind each soil site (i.e., there are five replicates for Atlas plants in Atlas soil and Temperate climate simulation).

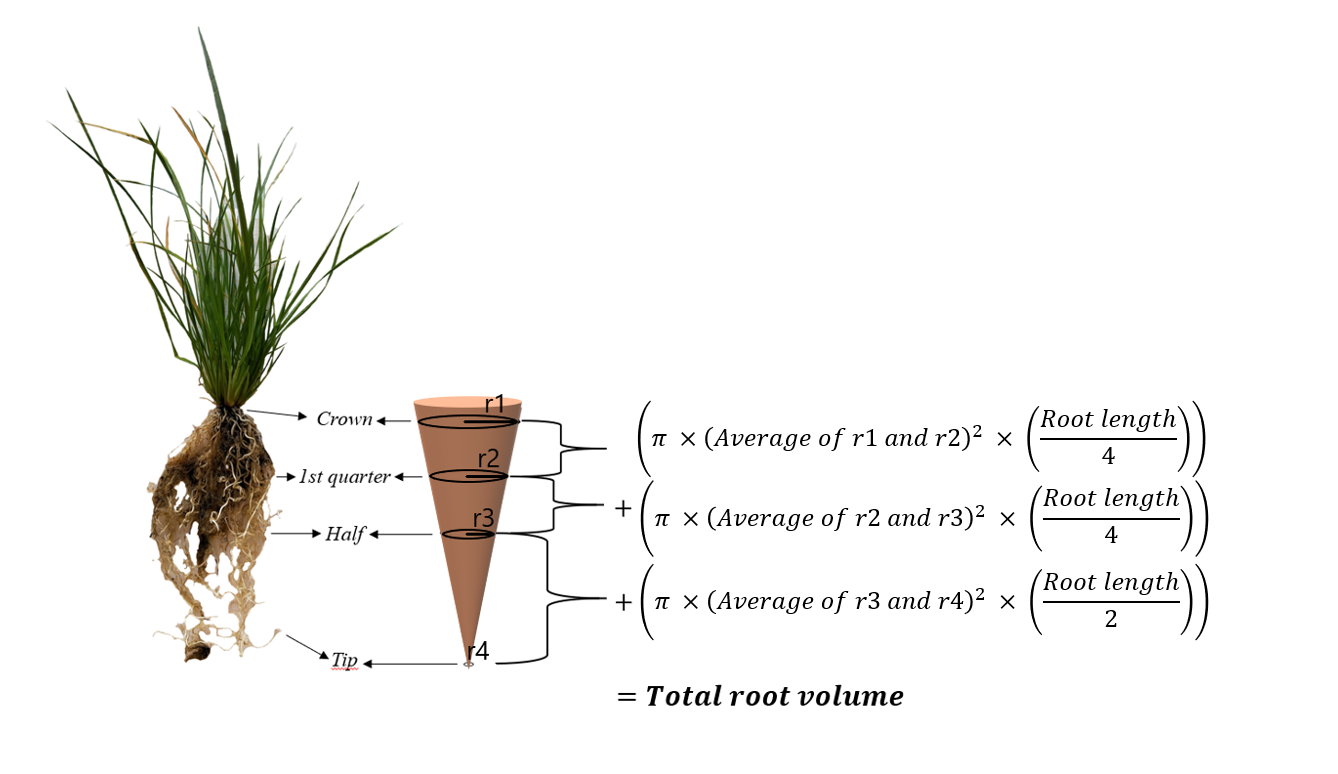

Figure S2: Root volume calculations. The radius of the crown and the radius of the root at the 1st quarter down its full length was averaged and used for the calculation of volume of the upper ¼ of the total root length. Average of 1st quarter and half is the mean of the radiuses measured on the first ¼ of total root length and in the middle of total root length. Average of half and tip is the mean of the radiuses measured in the middle of total root length and at the tip of the longest root. Root length is the total length from crown to the tip of the longest root when the roots are physically stretched out.

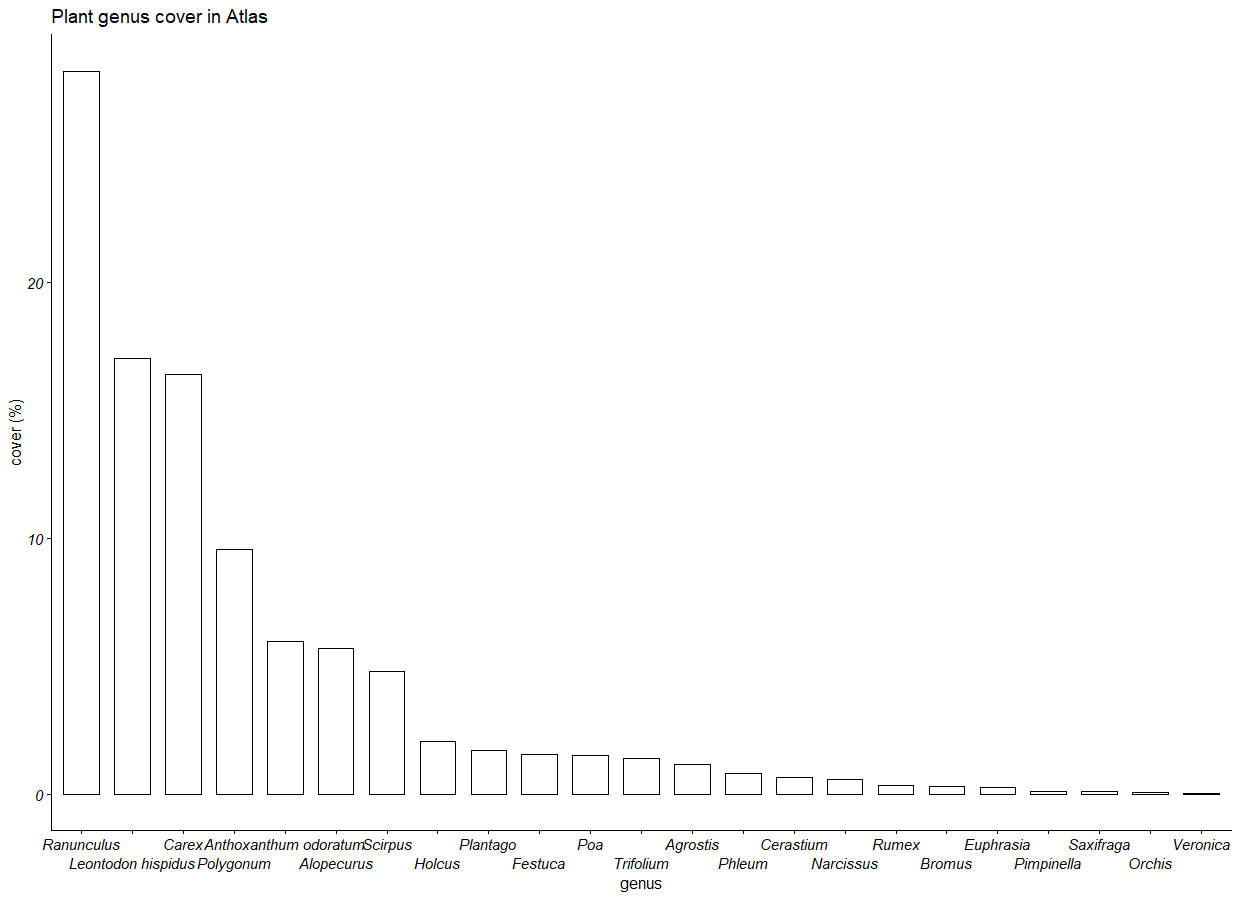

Figure S3: Plant genus cover in soil sampling site Atlas (Oukaïmeden). Cover represents relative cover of each genus in percentage.

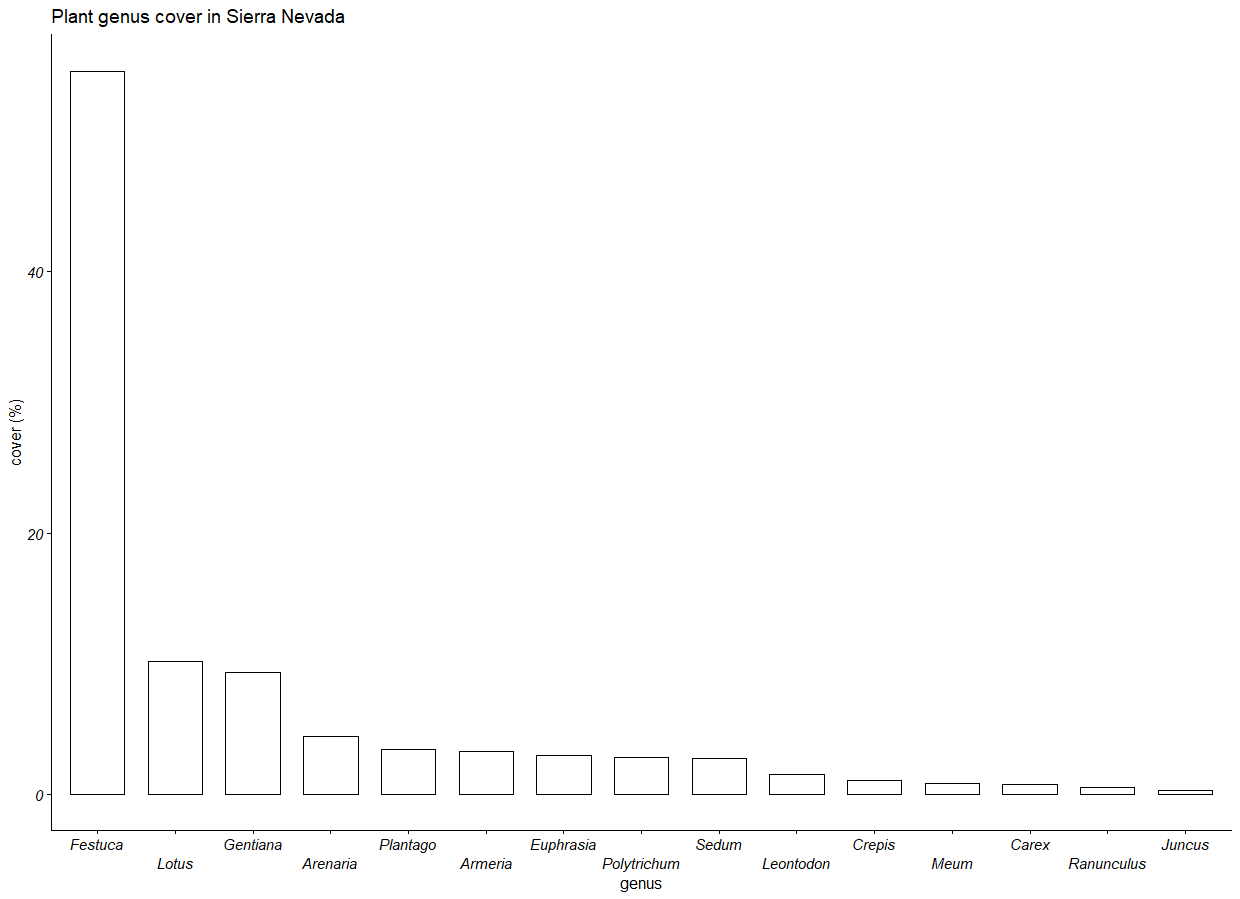

Figure S4: Plant genus cover in soil sampling site Sierra Nevada (Borreguiles). Cover represents relative cover of each genus in percentage.

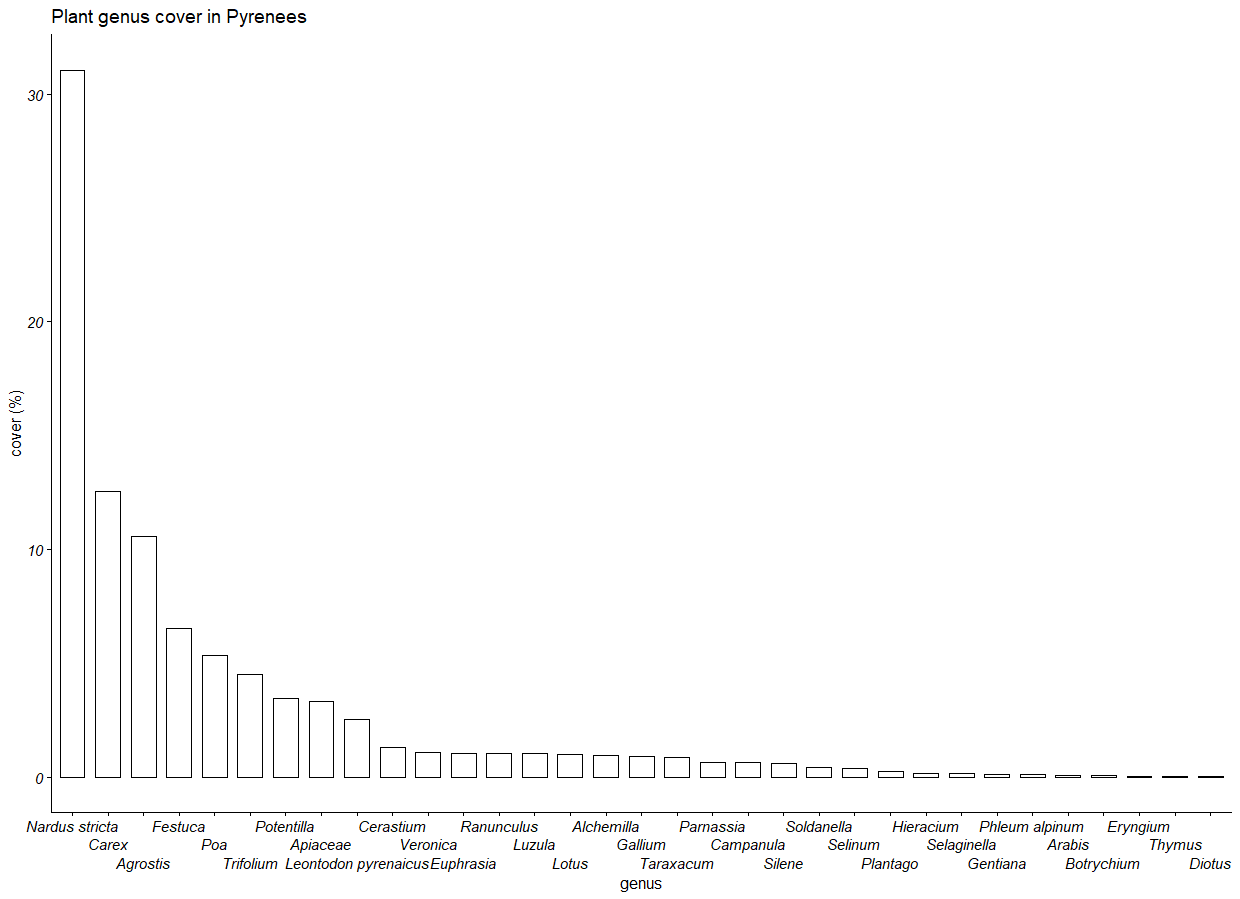

Figure S5: Plant genus cover in soil sampling site Pyrenees (Néouvielle). Cover represents relative cover of each genus in percentage.

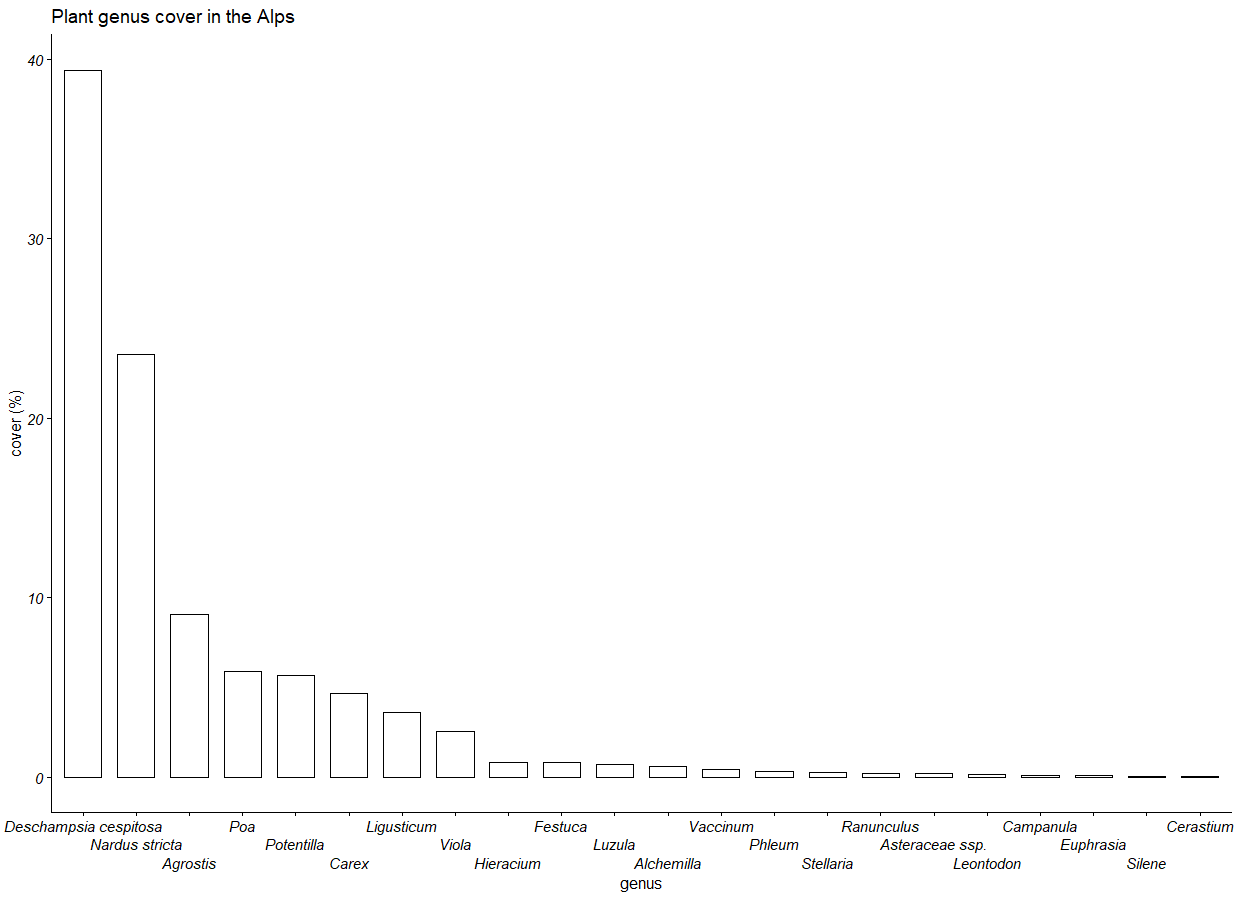

Figure S6: Plant genus cover in soil sampling site Alps (Flüela Pass). Cover represents relative cover of each genus in percentage.

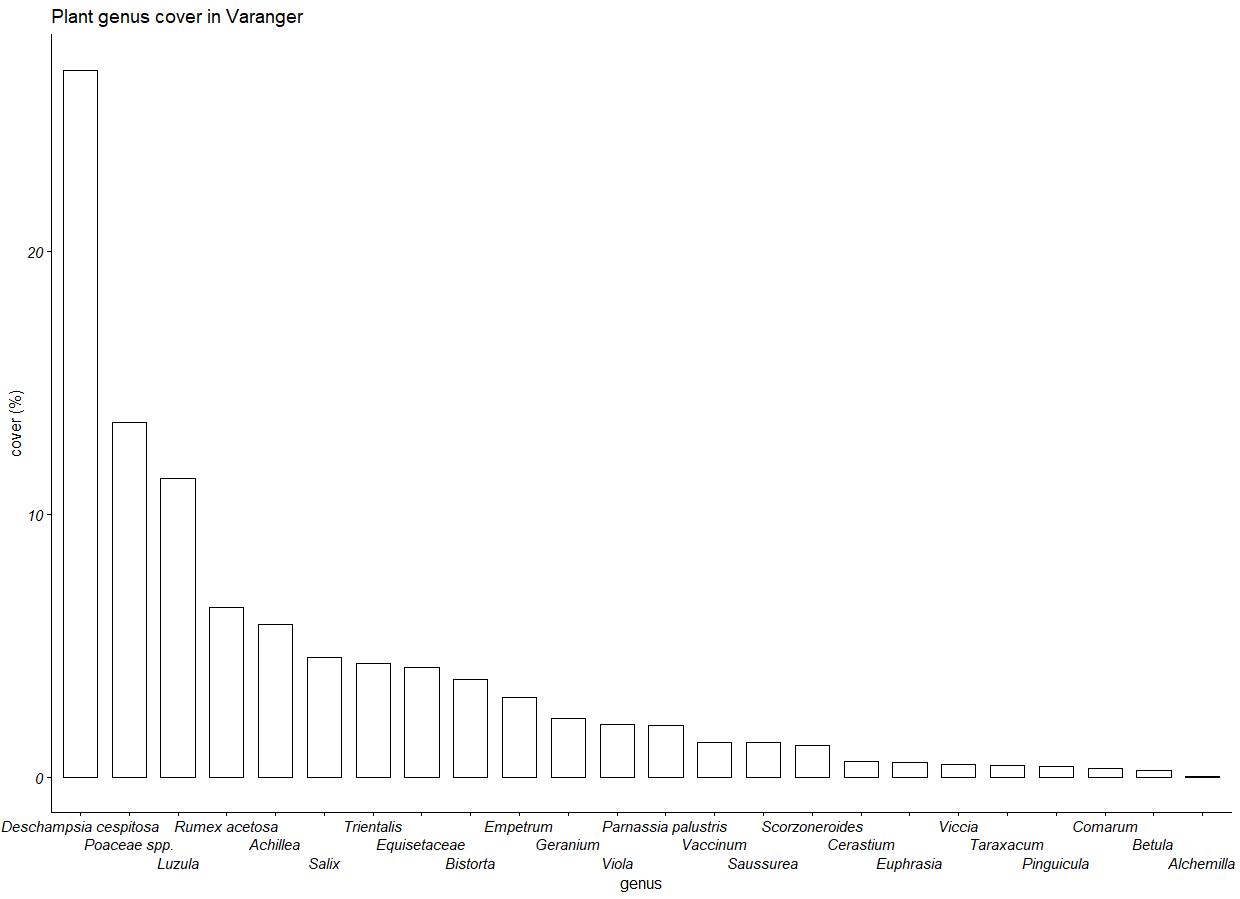

Figure S7: Plant genus cover in soil sampling site Varanger (Risfjorddalen, Store Salttjern, Kibergselva, Vesterelva, Finnvika, Sandfjordelva). Cover represents relative cover of each genus in percentage.

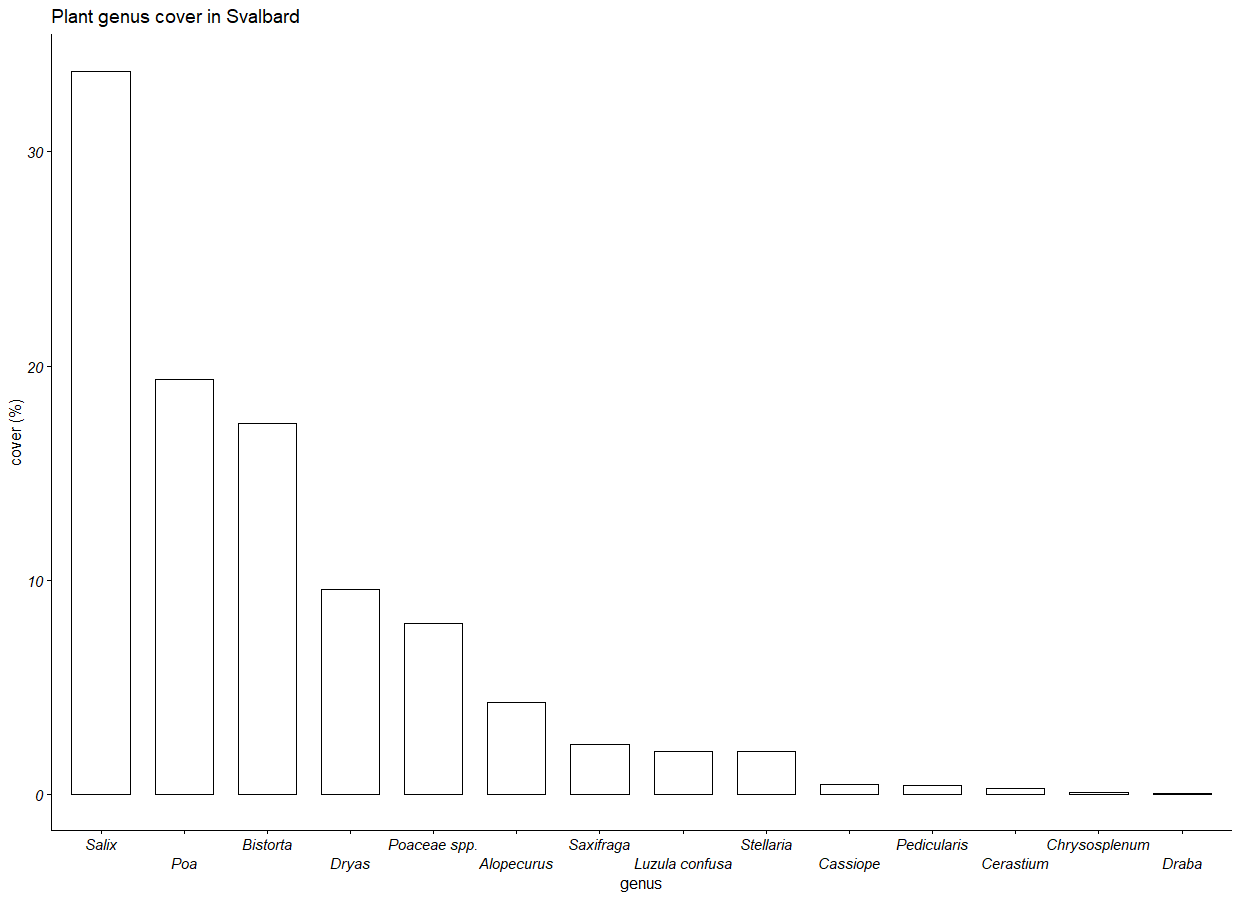

Figure S8: Genus cover in soil sampling site Svalbard (Adventdalen). Cover represents relative cover of each genus in percentage.

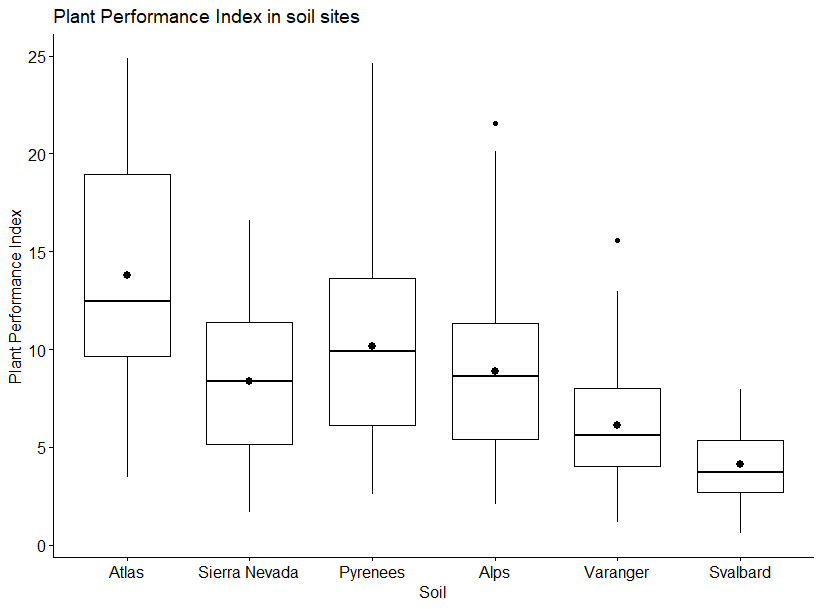

Figure S9: Plant performance index (PPI) from all plants in soil from each of the six sampling sites. Each box shows the mean (large black dot), median (middle horizontal line) and the lower and upper quartiles. Whiskers span the range of non-extreme values (less than 1.5 times the interquartile range), and the small dots show the outliers.

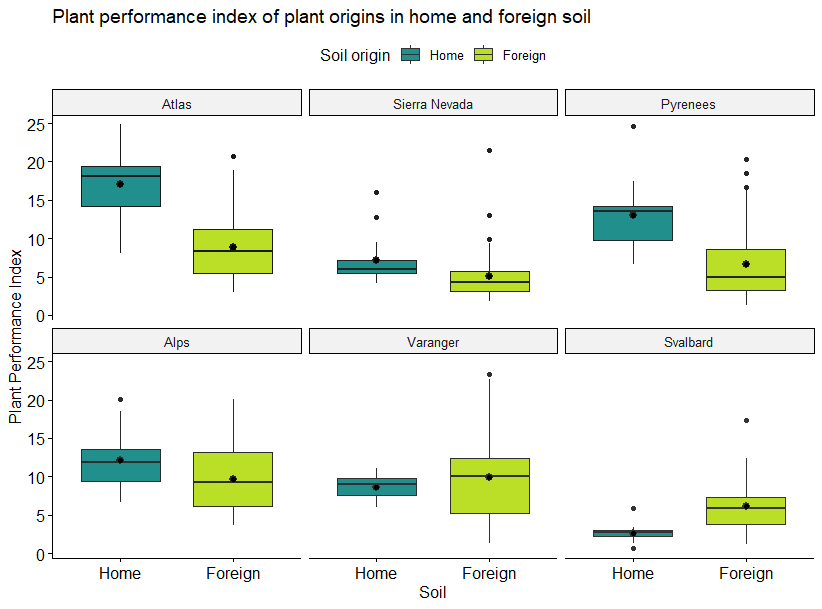

Figure S10: PPI in home and foreign soil facetted by plant origin. Each box shows the mean (large black dot), median (middle horizontal line) and the lower and upper quartiles. Whiskers span the range of non-extreme values (less than 1.5 times the interquartile range), and the small dots show the outliers. Boxes displaying plants grown in their home soil are coloured blue, whereas boxes displaying plants grown in foreign soil are coloured green.

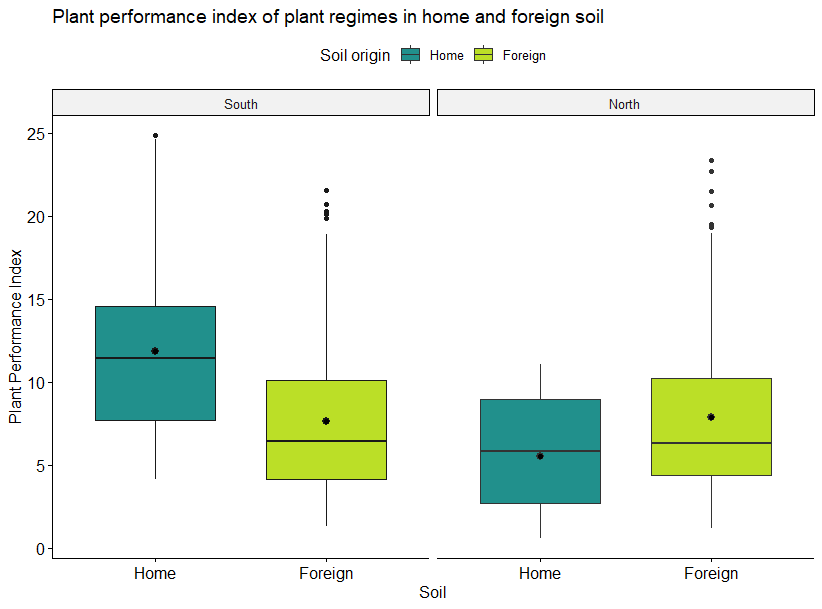

Figure S11: PPI in home and foreign soil for plants from all sites, separately. Each box shows the mean (large black dot), median (middle horizontal line) and the lower and upper quartiles. Whiskers span the range of non-extreme values (less than 1.5 times the interquartile range), and the small dots show the outliers. Boxes displaying plants grown in their home soil are coloured blue, whereas boxes displaying plants grown in foreign soil are coloured green.

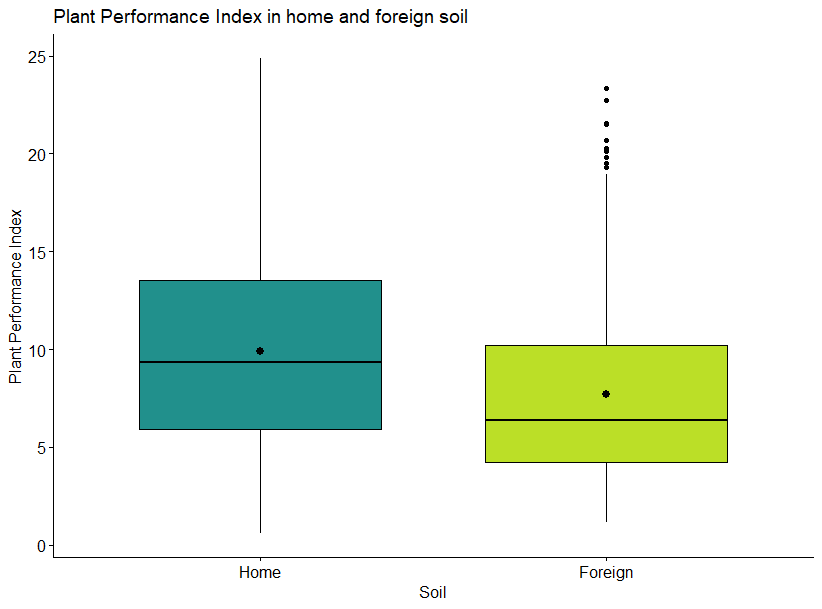

Figure S12: PPI in home and foreign soil for plants from all sites. Each box shows the mean (large black dot), median (middle horizontal line) and the lower and upper quartiles. Whiskers span the range of non-extreme values (less than 1.5 times the interquartile range), and the small dots show the outliers. Boxes displaying plants grown in their home soil are coloured blue, whereas boxes displaying plants grown in foreign soil are coloured green.

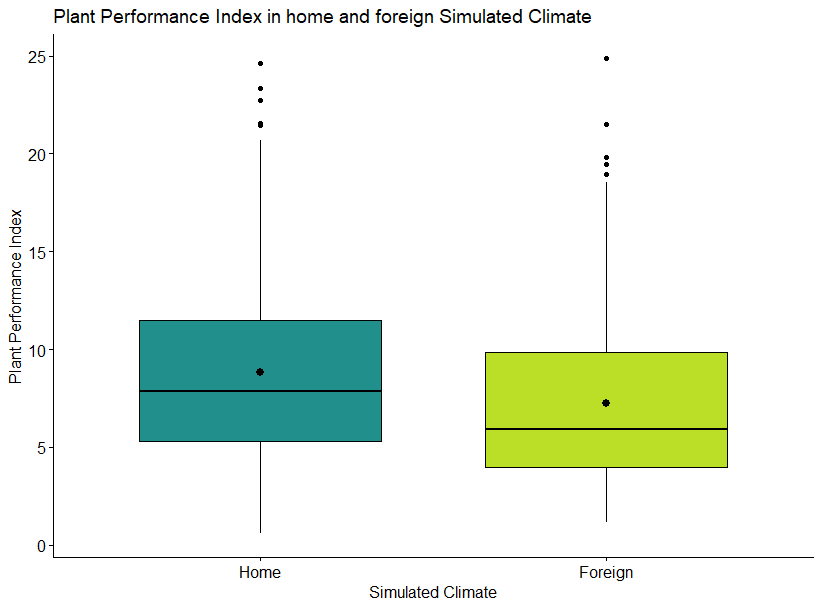

Figure S13: PPI in home and foreign simulated climate for plants from all sites. Each box shows the mean (large black dot), median (middle horizontal line) and the lower and upper quartiles. Whiskers span the range of non-extreme values (less than 1.5 times the interquartile range), and the small dots show the outliers. Boxes displaying plants grown in their home soil are coloured blue, whereas boxes displaying plants grown in foreign soil are coloured green.

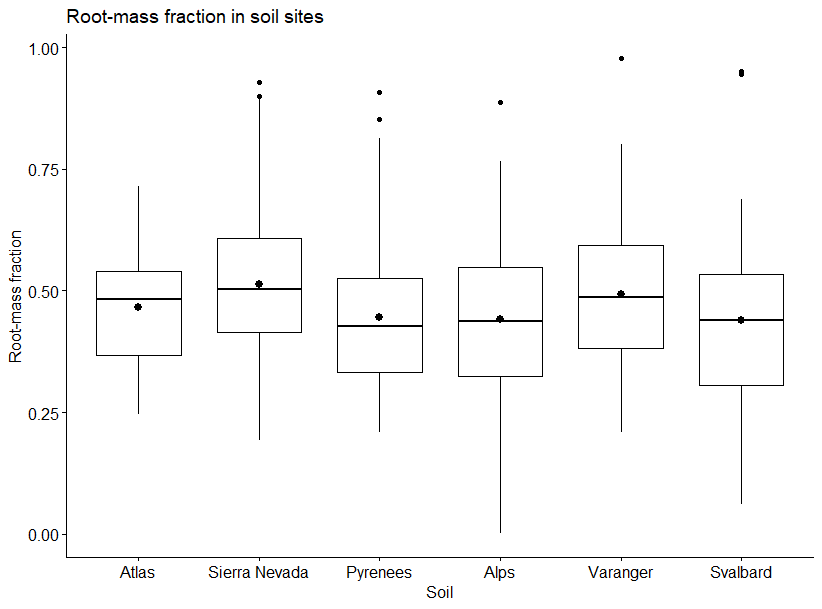

Figure S14: Root-mass fraction (RMF) from all plants in soil from each of the six sampling sites. Each box shows the mean (large black dot), median (middle horizontal line) and the lower and upper quartiles. Whiskers span the range of non-extreme values (less than 1.5 times the interquartile range), and the small dots show the outliers

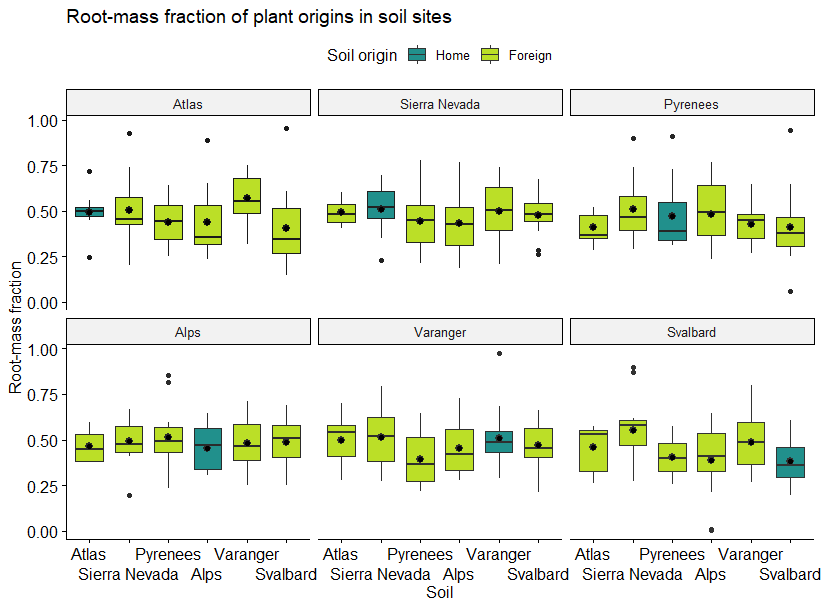

Figure S15: Root -mass fraction (RMF) in soils of different soil site origins, facetted by plant origin. Each box shows the mean (large black dot), median (middle horizontal line) and the lower and upper quartiles. Whiskers span the range of non-extreme values (less than 1.5 times the interquartile range), and the small dots show the outliers. Boxes displaying plants grown in their home soil are coloured blue, whereas boxes displaying plants grown in foreign soil are coloured green.

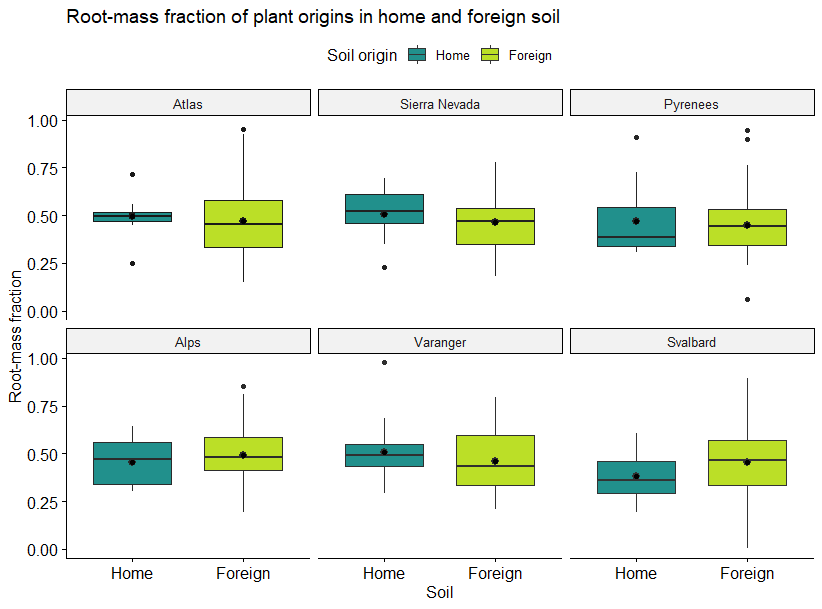

Figure S16: Root-mass fraction in home and foreign soil, facetted by plant origin. Each box shows the mean (large black dot), median (middle horizontal line) and the lower and upper quartiles. Whiskers span the range of non-extreme values (less than 1.5 times the interquartile range), and the small dots show the outliers. Boxes displaying plants grown in their home soil are coloured blue, whereas boxes displaying plants grown in foreign soil are coloured green.

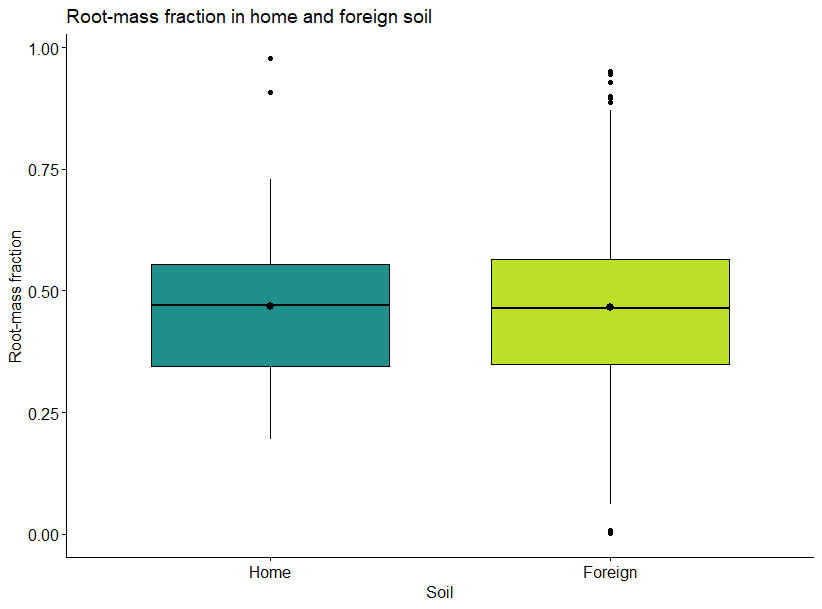

Figure S17: Root-mass fraction in home and foreign soil for plants from all sites. Each box shows the mean (large black dot), median (middle horizontal line) and the lower and upper quartiles. Whiskers span the range of non-extreme values (less than 1.5 times the interquartile range), and the small dots show the outliers. Boxes displaying plants grown in their home soil are coloured blue, whereas boxes displaying plants grown in foreign soil are coloured green.

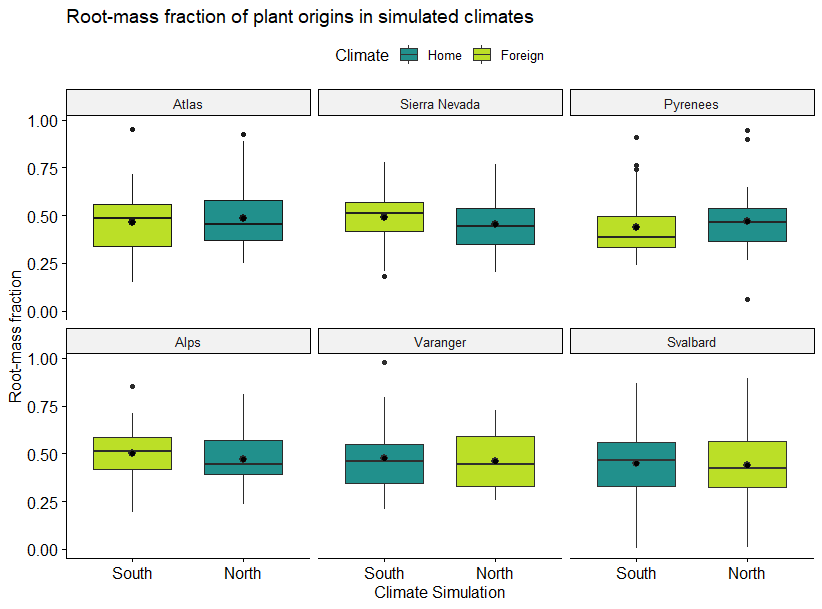

Figure S18: Root-mass fraction in the two simulated climates, facetted by plant origin. Each box shows the mean (large black dot), median (middle horizontal line) and the lower and upper quartiles. Whiskers span the range of non-extreme values (less than 1.5 times the interquartile range), and the small dots show the outliers. Boxes displaying plants grown in their home soil are coloured blue, whereas boxes displaying plants grown in foreign soil are coloured green.

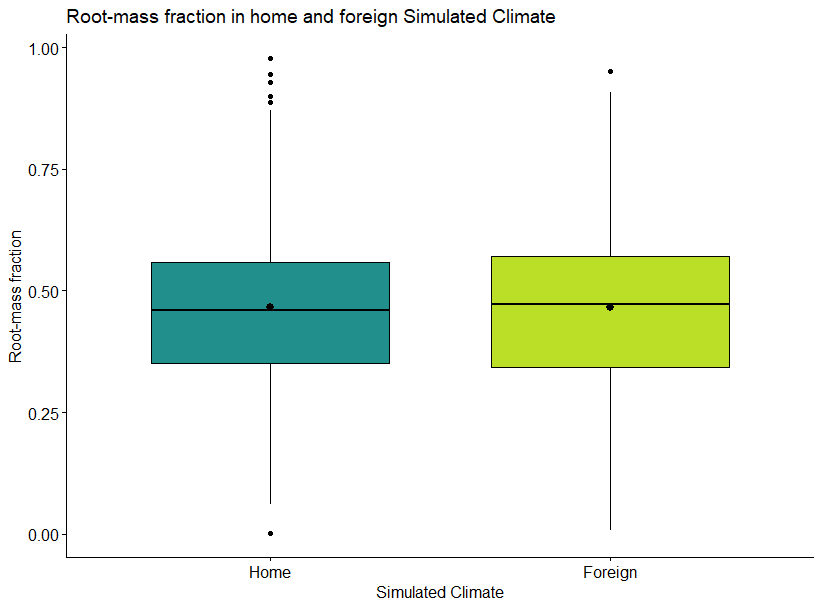

Figure S19: Root-mass fraction in home and foreign simulated climate for plants from all sites. Each box shows the mean (larger black dot), median (middle horizontal line) and the lower and upper quartiles. Whiskers span the range of non-extreme values (less than 1.5 times the interquartile range), and the small dots show the outliers. Boxes displaying plants grown in their home soil are coloured blue, whereas boxes displaying plants grown in foreign soil are coloured green.

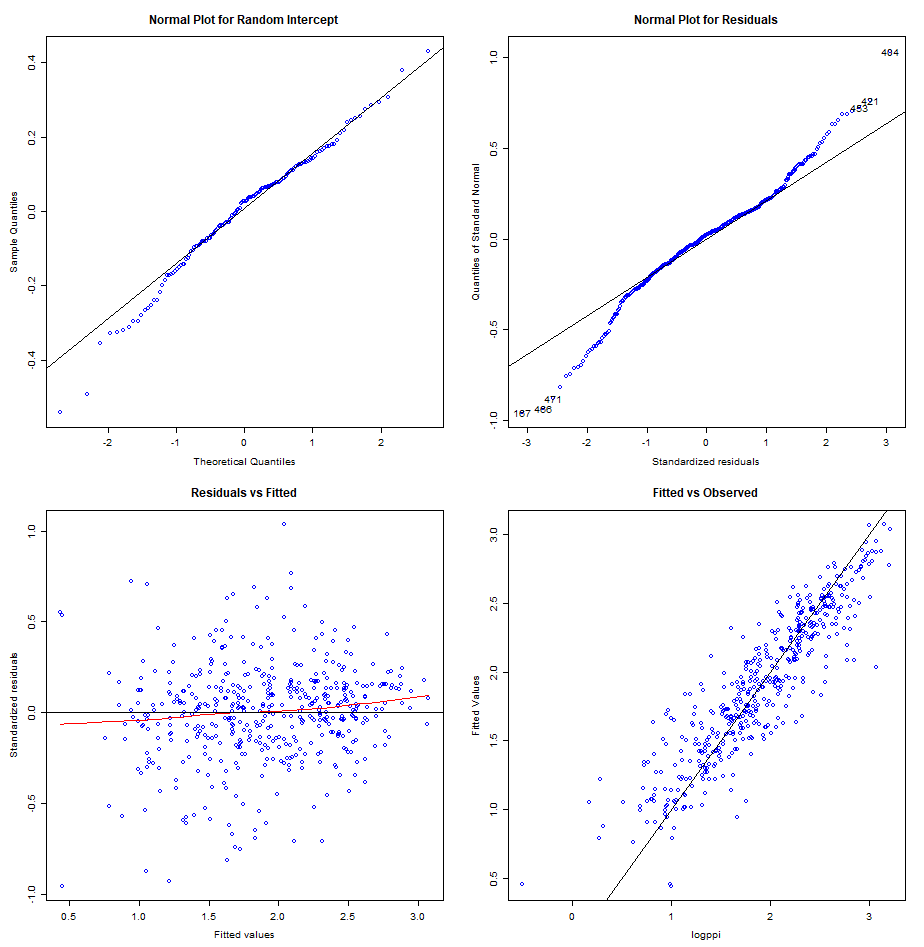

Figure S20: Residual plots for the linear model with PPI as response variable using log transformation after removal of superfluous predictors.
